# On the discovery of ADRAM, an experience-dependent long noncoding RNA that drives fear extinction through a direct interaction with the chaperone protein 14-3-3

**DOI:** 10.1101/2021.08.01.454607

**Authors:** Xiang Li, Qiongyi Zhao, Ziqi Wang, Wei-Siang Liau, Dean Basic, Haobin Ren, Paul R. Marshall, Esmi L. Zajaczkowski, Laura J. Leighton, Sachithrani U. Madugalle, Mason Musgrove, Ambika Periyakaruppiah, Jichun Shi, Jianjian Zhang, John S. Mattick, Timothy R. Mercer, Wei Wei, Timothy W. Bredy

## Abstract

Long-noncoding RNA (lncRNA) comprise a new class of genes that have been assigned key roles in development and disease. Many lncRNAs are specifically transcribed in the brain where they regulate the expression of protein-coding genes that underpin neuronal function; however, their role in learning and memory remains largely unexplored. We used RNA Capture-Seq to identify a large population of lncRNAs that are expressed in the infralimbic cortex of adult male mice in response to fear-related learning, with 14.5% of these annotated in the GENCODE database as lncRNAs with no known function. We combined these data with cell-type-specific ATAC-seq on neurons that had been selectively activated by fear-extinction learning, and revealed 434 lncRNAs derived from enhancer regions in the vicinity of protein-coding genes. In particular, we discovered an experience-induced lncRNA called ADRAM that acts as both a scaffold and a combinatorial guide to recruit the brain-enriched chaperone protein 14-3-3 to the promoter of the memory-associated immediate early gene Nr4a2. This leads to the expulsion of histone deactylases 3 and 4, and the recruitment of the histone acetyltransferase creb binding protein, which drives learning-induced Nr4a2 expression. Knockdown of ADRAM disrupts this interaction, blocks the expression of Nr4a2, and ultimately impairs the formation of fear-extinction memory. This study expands the lexicon of experience-dependent lncRNA activity in the brain, highlights enhancer-derived RNAs (eRNAs) as key players in the epigenetic regulation of gene expression associated with fear extinction, and suggests eRNAs, such as ADRAM, may constitute viable targets in developing novel treatments for fear-related anxiety disorders.

## INTRODUCTION

The extinction of conditioned fear, the reduction in responding to a feared cue, which occurs when the cue is repeatedly presented without any adverse consequence, is an evolutionarily conserved behavioural adaptation that is critical for survival. Efforts to understand the mechanisms of this important form of learning have increased recently as it is a preclinical model for the treatment of anxiety disorders. Like other forms of learning, long-lasting memory for fear extinction depends on coordinated changes in gene expression, particularly in the infralimbic prefrontal cortex (ILPFC) (Martin et al., 2000; Bruel-Jungerman et al., 2007; Alberini, 2009). In recent years, we and others have shown that this process involves a tightly controlled interplay between transcriptional machinery and epigenetic mechanisms (reviewed in Marshall and Bredy, 2020). Indeed, a wide variety of chromatin and DNA modifications have been shown to play an essential role in various forms of learning and the establishment of long-term memory (Bredy et al., 2007, Vescey et al, 2007; Wei et al., 2012; Gräff et al., 2014; Li et al., 2014, 2019; Feng et al, 2015; Lepack et al, 2020).

Although much progress has been made in understanding how epigenetic modifiers are directed to their requisite sites of action across the genome during early development, it remains to be fully determined how this specificity is conferred in the adult brain, particularly within the context of learning and memory. Long noncoding RNAs (lncRNAs) comprise a class of genes that have recently gained attention as important regulators of cellular function due to their multidimensional capacity to function as decoys for transcription factors, as guides to direct chromatin modifiers, or as modular scaffolds in the nucleus (Mercer and Mattick, 2013). LncRNAs, defined as any RNA longer than 200nt and without protein coding potential, are expressed in a highly cell-type and spatiotemporally specific manner in the adult brain (Mercer et al, 2008), with 40% of all lncRNAs identified to date shown to be enriched in neurons. It has therefore been proposed that lncRNAs are uniquely positioned to mediate rapid responses to environmental stimuli and to promote cognition (Spadaro and Bredy, 2012; Liau et al., 2021). In agreement with this idea, the nuclear-enriched lncRNA *Gm12371* influences hippocampal dendritic morphology and synaptic plasticity (Raveendra et al., 2018), the nuclear antisense lncRNA *AtLAS* regulates synapsin II polyadenylation and AMPA receptor trafficking (Ma et al., 2020), and an association between the lncRNA *LONA* and synaptic plasicity and spatial memory (Li et al., 2018) has been reported. Not surprisingly, lncRNAs have also been implicated in the regulation of gene expression underlying neuropsychiatric disorders characterised by impaired cognition, including drug addiction, depression, impulsivity, schizophrenia, and anxiety (Barry et al., 2014; Spadaro et al., 2015; Xu et al., 2020; Issler et al., 2020; Labonte et al., 2020).

Given the increasing recognition that lncRNAs play important roles in brain function and neuropsychiatric disease, we considered their impact on fear extinction. We first used targeted RNA sequencing to reveal lncRNAs that were induced in response to fear-related learning and its extinction. The complex isoform architecture of the newly identified lncRNAs was then resolved by ATAC-sequencing, with chromatin and RNA immunoprecipitation analysis being employed to functionally characterize the lncRNA-mediated epigenetic regulation of target gene expression. Finally, lentiviral-mediated knockdown and antisense oligonucleotide injections were used to investigate the causal mechanisms by which ADRAM, a novel lncRNA derived from a proximal enhancer, regulates the expression of the immediate early gene Nr4a2 and drives the formation of fear extinction memory.

## RESULTS

### A substantial number of lncRNAs are expressed in the ILPFC in response to fear-related learning

Most lncRNAs are expressed in low abundance and with cell-type specificity. As a result, they are often missed during the analysis of complex tissues such as the brain (Deveson et al., 2017). We therefore adopted a method called RNA capture sequencing (Capture-Seq), which provides the additional sensitivity required to determine the full repertoire of learning-induced lncRNAs in the adult ILPFC. RNA Capture-seq uses tiling oligonucleotides to enrich for RNA targets of interest prior to sequencing, resulting in a dramatic increase in the sensitivity to detect rare transcripts (Mercer et al., 2011; 2014). In the following experiment, a panel targeting 190,689 probes comprising 28,228 known and predicted mouse lncRNAs, which was previously developed to improve the annotation of brain-enriched lncRNA (Bussotti et al, 2016), was used to identify lncRNAs in the ILPFC that are expressed in response to fear learning and during the formation of fear-extinction memory. We trained mice using a standard cued fear-conditioning task followed by either novel context exposure (retention control, RC) or extinction training (EXT) and, immediately after training, the ILPFC was extracted and RNA prepared for downstream analysis.

Using RNA Capture-seq, with an average of 68 million (83%) uniquely mapped reads per pooled library (**Supplemental Table 1**), we identified a total of 23,514 lncRNAs that were expressed following RC and EXT, with many being novel (66%, 15,439) or listed in the GENCODE database as transcripts (14.5%, 3,412) of unknown function. *Meg3* (Chandra et al., 2018), *Malat1* (Wu et al., 2018), and *Gomafu* (Spadaro et al., 2015) were among the most abundantly expressed brain-enriched lncRNAs that have been functionally characterized, although no differential expression between RC and EXT mice was detected (**Supplemental Table 2**). A transcript-level expression analysis comparing the RC and EXT groups revealed that none of the detected lncRNAs reached the threshold (FDR<0.05) to be considered differentially expressed following learning. This is perhaps surprising given the historical reliance on concluding that a gene is relevant or causal based on whether it is up- or down-regulated. However, emerging evidence indicates that the apparently low levels of expression of lncRNAs generally reflects their high cell-type specificity (Mercer et al., 2008; Cabili et al., 2015; Seiler et al., 2017; Deveson et al., 2017) and there are increasing examples of lncRNAs with highly restricted spatial expression that impact brain development and function (Cajigas et al., 2018; Hollensen et al., 2020; Perry et al., 2018; Raveendra et al., 2018; Wang et al., 2021; Grinman et al., 2021).

LncRNAs exhibit extreme variations in length, have a modular architecture, and undergo complex patterns of alternative splicing (Deveson et al., 2018), all of which can impact their functional activity independent of their level of expression. It is well established that lncRNAs exert their regulatory influence in a context-and state-dependent manner that is contingent on the cellular compartment in which they are expressed. In support of this idea, we previously observed this to be the case with the lncRNA *Gomafu (Miat),* which was observed to function both *in cis* as a scaffold for the polycomb complex within the local genome environmnent (Spadaro et al., 2015), and *in trans* as a decoy for splicing factors QK1 and SRSF1 in the nucleolus (Barry et al., 2014). Moreover, although *Malat1* is one of the most highly abundant lncRNAs to be identified to date, its structure state is heavily influenced by RNA modification, which alters its ability to interact with specific RNA binding proteins, thereby determining its functional state independent of transcript abundance (Liu et al., 2017). In addition, the lncRNA *Neat1* has a highly complex modular structure, which confers its ability to act as a scaffold in the assembly of paraspeckles through phase separation (Yamazaki et al., 2018). This may influence how it participates in behavioural responses to stress (Kucharski et al., 2020) and as an architect of chromatin modification supporting age-related spatial memory processes (Butler et al., 2019). It is increasingly becoming evident that there are many factors beyond transcriptional abundance that determine whether a lncRNA is functionally relevant (Liau et al., 2021). We therefore decided to look more deeply into *how* lncRNAs are regulated in the ILPFC, and *how* they are functionally activated in response to fear extinction learning.

### LncRNAs exhibit unique, context-dependent, chromatin accessibility profiles

LncRNAs are frequently found antisense, bidirectional or in close proximity to key protein coding-genes. Furthermore, many lncRNAs mediate their actions *in cis* by regulating the local chromatin context of neighbouring protein-coding genes. Based on this, we next analysed the genomic organisation of lncRNAs in cells that have been activated by learning. In an independent cohort of RC and EXT trained mice, the activity-regulated cytoskeleton-associated protein (*Arc*) and the neuronal nuclear marker *NeuN* was used to tag specific populations of neurons that had been selectively activated by RC or EXT training, which were then isolated by fluorescence-activated cell sorting (FACS) and prepared for downstream analysis (Li et al., 2019; **Supplemental Figure 1**). ATAC-seq (assay for transposase-accessible chromatin using sequencing) was employed on Arc+ neuronal populations in order to resolve the genome-wide landscape of chromatin accessibility following RC and EXT learning in activated derived from the ILPFC. Overall, with an average of 86 million (86%) uniquely mapped reads per pooled library, we found that chromatin accessible regions exhibited a similar genomic distribution in both conditions (**Supplemental Table1, Supplemental Figure 2**).

Focusing specifically on the Arc+ neurons of EXT mice, we found that accessible chromatin regions were more likely to be enriched at the 5’ UTR (17.5%) rather than the 3’ (2.5%) UTR, which occurred with a similar frequency to transcription start sites (16.8%) (**Figure 1A**). A significant proportion of ATAC peaks were found in intronic (43%) and intergenic (27.3%) regions, indicating a potentially rich source of active genomic regions from which many novel lncRNAs may be derived (**Figure 1A**). Indeed, the majority of expressed lncRNAs were found in either intragenic (11,496: 49% of total) or intergenic regions distal (>10kb) to protein-coding genes (9,909: 42% of total). In addition, we found 2,109 lncRNAs (∼9% of total) were expressed in genomic regions in the vicinity (<10kb) of protein-coding genes. Among these 2,109 proximal lncRNAs, 42% (892) were associated with open chromatin, which was markedly higher than the proportion of intergenic (16%) or intragenic (26%) lncRNAs (**Supplemental Table 2**).

**Figure 1.**
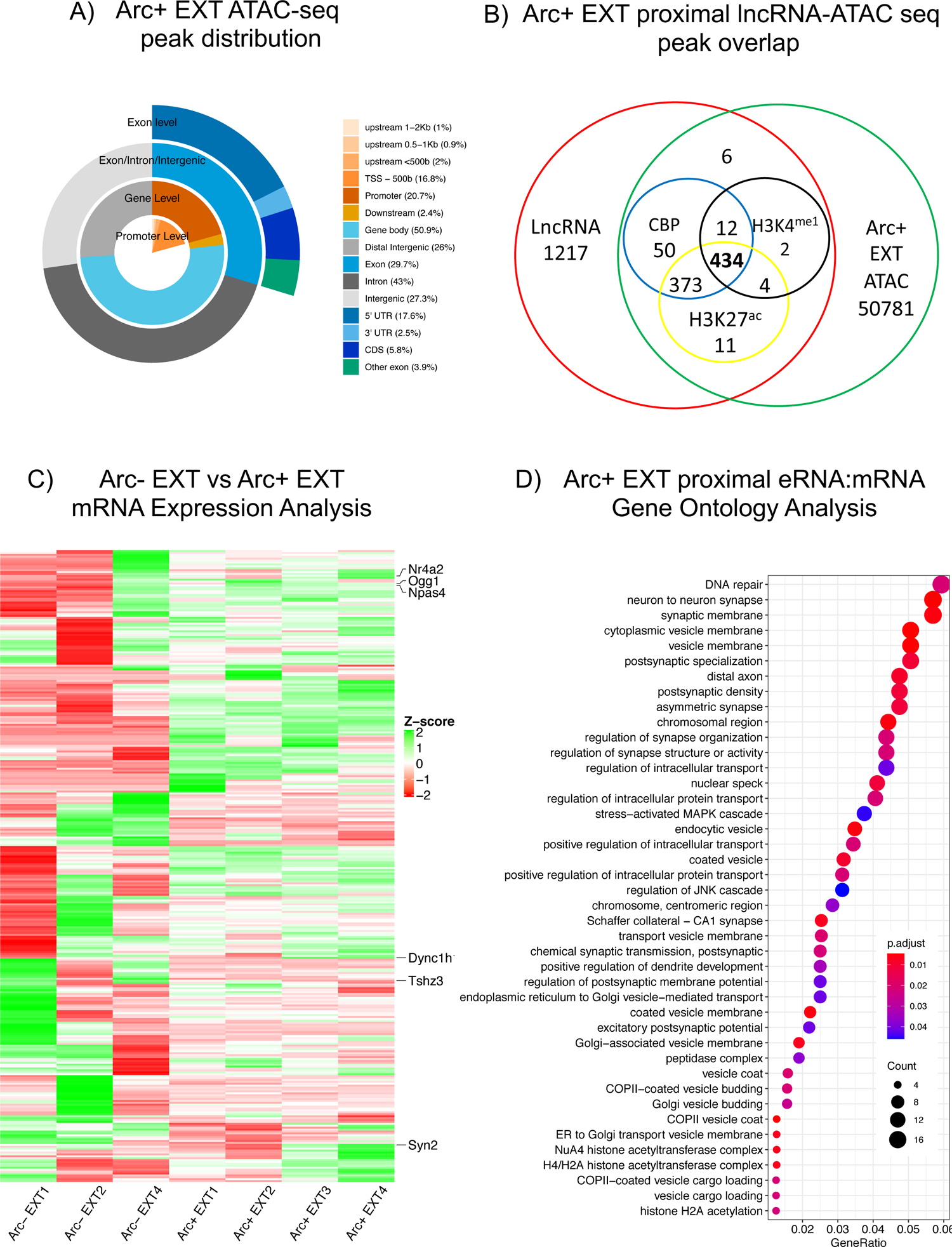
LncRNAs derived from enhancer elements are induced by fear-extinction learning and correlate with proximal protein-coding gene expression in the adult prefrontal cortex. A) Genomic distribution of ATAC peaks in Arc+ neurons that have been activated by fear-extinction learning. B) The Venn diagram highlights 434 proximal lncRNAs overlapping with lncRNA capture-seq, Arc+EXT ATAC-seq, as well as H3K27^ac^, CBP and H3K4^me1^ enriched genomic regions. C) Heatmap of eRNA-associated mRNA expression in quiescent (ARC-EXT) versus activated (ARC+ EXT) neurons (n=3 biological replicates for ARC-EXT; n=4 biological replicates ARC+ EXT; Red: decreased expression and Green: increase expression). D) Gene ontology (GO) analysis for proximal protein-coding genes located <10 kb downstream of the 434 eRNA loci. Top 30 significantly enriched GO terms are shown in the dot plot.

Since lncRNAs are expressed in a highly cell-type and spatiotemporally specific manner, we next considered the mechanisms underlying the cell specific and state-dependent expression of learning-induced lncRNAs. Open chromatin-accessibility sites were compared with publicly available data on H3K27^ac^, H3K4^me1^ and CBP occupancy in the adult brain that occurs following behavioural training (Halder et al., 2016). In Arc+ RC mice, amongst the 60,962 significant chromatin-accessibility sites, we found that 25.2% (5,915) of dynamically-expressed lncRNAs were associated with open chromatin and 2,718 regions exhibited features of active enhancers in response to fear conditioning (**Supplemental Figure 1C**). In the Arc+ neurons of EXT mice, a total of 51673 peaks was detected by ATAC-seq, with 23.2% (5,460) of all lncRNAs being associated with increased chromatin accessibility. We found 2,501 regions exhibiting features of active enhancers in response to fear extinction training (**Supplemental Table 2**). Together, these data show that the active regulation of numerous lncRNAs upon learning is associated with markers of enhancer activity, and suggest that the experience-dependent expression of lncRNA in the adult brain may be more widespread than currently appreciated.

### A significant population of enhancer-derived lncRNAs that are induced by fear extinction learning positively correlate with proximal protein-coding gene expression

We next focused on lncRNAs that were associated with enhancers (termed eRNAs) using ATAC-seq as well as the publicly available CBP, H3K27^ac^ and H3K4^me1^ signatures. We identified 434 putative eRNAs in Arc+ EXT neurons that overlapped with all four enhancer markers (**Figure 1B**). We found that in mRNAs positioned downstream of these eRNAs, there was a tighter correlation in overall expression in Arc+ EXT activated neurons than in quiescent Arc-EXT neurons derived from the same brain region (**Figure 1C**), suggesting that proximal eRNA activity may serve to synchronize patterns of mRNA expression in response to fear extinction learning. A gene ontology (GO) analysis was then performed to explore the putative functional networks of protein-coding genes positioned proximal to eRNAs expressed during fear extinction learning. We found that these genes were most significantly enriched for GOs linked to ‘cytoplasmic vesicle membrane’, ‘neuron to neuron synapse’, ‘vesicle membrane’ and ‘synaptic membrane’ (**Figure 1D and Supplemental Table 3**). Other significant GO terms that contained fewer transcripts but were nonetheless interesting in the context of fear extinction were ‘DNA repair’, ‘nuclear speckles’ and several vesicle-related or postsynaptic-related clusters.

Next, 10 candidate proximal lncRNA:mRNA pairs were selected for validation based on the diversity of their genomic organization, their variable length (600-3000nt), and the fact that their associated protein-coding genes have previously been shown to be involved in neuronal plasticity and/or memory-related processes. Examples of these include two known lncRNAs with bidirectional promoters (*Gm26559:Map1b, Gm17733:Syn2),* another known intergenic lncRNA (*A730063M14Rik:Agap2),* an antisense lncRNA *(Gm15492:Ogg1),* and two functionally uncharacterised transcripts *(Gm13830:Srsf9 and BB557941:Nr4a2)*. In each case, we observed a strong positive correlation between the expression of the lncRNA and its proximal protein-coding mRNA, as determined by qPCR (**Figure 2A-F, and Supplemental Figure 3**). The discrepancy between the lncRNA capture-seq results and the qPCR findings likely stems from the fact that qPCR focuses solely on a single exon whereas sequencing provides an average of read coverage across the entire transcript, which then undergoes a more stringent statistical test in order to counteract the problem of multiple testing. Indeed, an examination of the read pile up across the ADRAM locus suggests there are more transcripts associated with the 3rd exon, which is specifically targeted by our PCR primer design. Taken together, our data suggest that there is a positive relationship between inducible eRNAs and downstream protein-coding RNA expression. Given that the immediate early gene *Nr4a2* is directly involved in learning and memory (McNulty et al., 2012; Chatterjee et al., 2020), we subsequently focused our investigation on the mechanistic relationship between its proximal lncRNA *BB557941* (**Figure 2G**), which we call *ADRAM* (**A**ctivity-**D**ependent lnc**R**NA **A**ssociated with **M**emory) and the epigenetic regulation of *Nr4a2,* as well as the role of this relationship in fear extinction.

**Figure 2.**
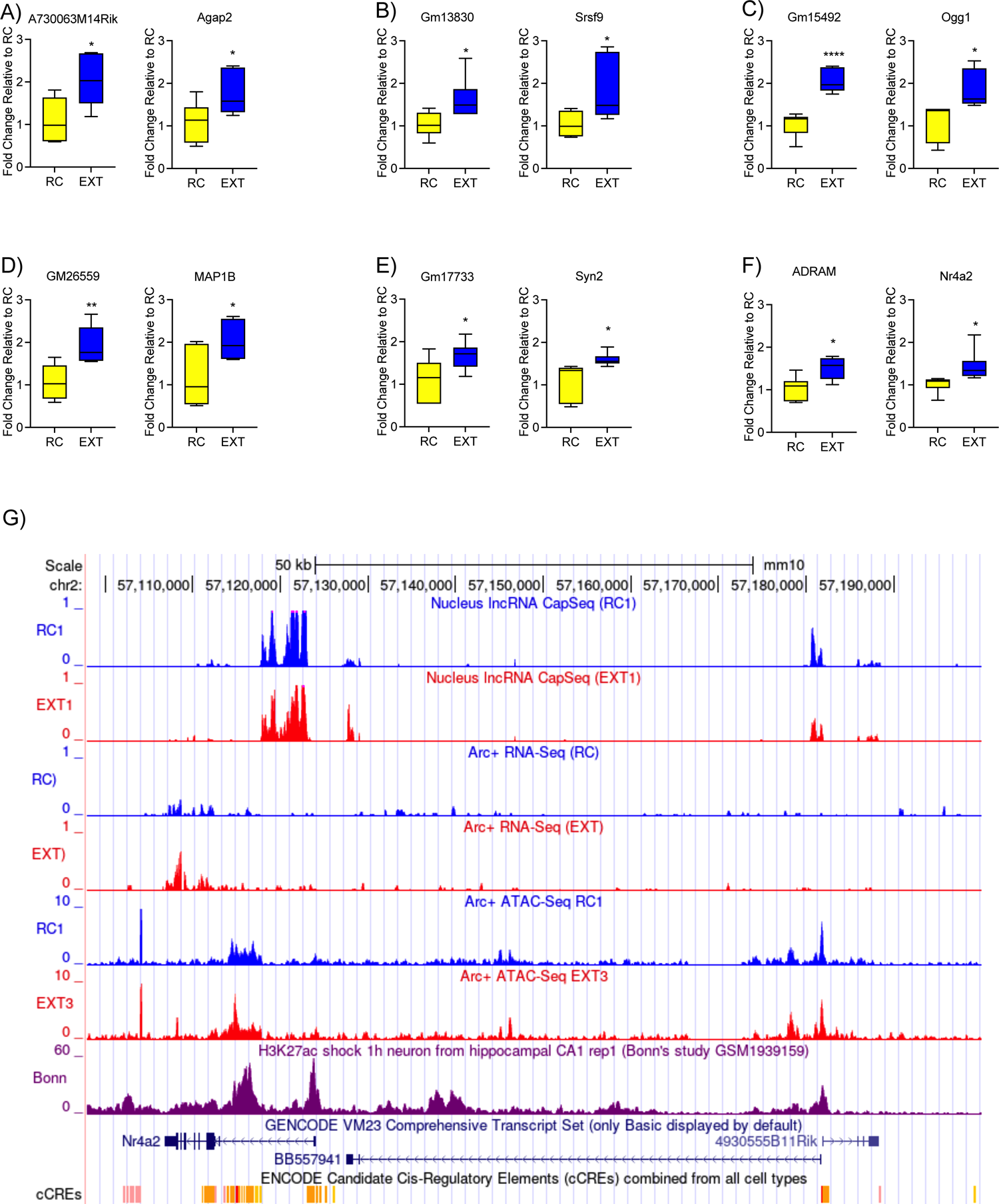
Extinction training leads to a correlated increase in the expression of proximal eRNAs and their downstream mRNAs. A) A730063M14Rik:Agap2, B) Gm17733:Syn2, C) Gm15492:Ogg1, D) Gm26559:Map1b, E) Gm13830:Srsf9, and F) BB557941:Nr4a2. (n=6 biological replicates per group, two-tailed unpaired Student’s t-test). Error bars represent S.E.M. *p<0.05, **p<0.01, ****p<0.0001. G) Representative UCSC genome browser track showing ADRAM, Nr4a2, and H3K27^ac^ peaks in RC and EXT trained mice.

### Learning-induced *ADRAM* expression is necessary for the induction of the immediate early gene *Nr4a2,* and is required for fear-extinction memory

Since *ADRAM* and *Nr4a2* are both upregulated in response to fear-extinction learning, we investigated whether *ADRAM* expression is necessary for the induction of the *Nr4a2*. First, using fluorescence in situ hybridization (FISH), we determined that ADRAM is selectively expressed in the nucleus of the cortical pyramidal neurons, *in vitro* (**Supplemental Figure 4**). Next, in order to elucidate its functional role in fear extinction, antisense oligonucleotides (ASOs) were used to knock down ADRAM expression, *in vivo*. Three different ASOs targeting *ADRAM* were first tested on primary cortical neurons, *in vitro* (**Supplemental Figure 5**). ASO1 exhibited the best knockdown efficiency (at 200nM). Injection of ASO1 into ILPFC neurons prior to fear-extinction training (**Supplemental Figure 6**) resulted in a significant reduction in learning-induced *ADRAM* expression (**Figure 3A**) and blocked the induction of *Nr4a2* mRNA expression (**Figure 3B**). Importantly, knocking down *ADRAM* prior to fear-extinction training (**Figure 3C**) had no effect on within-session performance during extinction training (**Figure 3D**), and there was also no effect of ADRAM knockdown on preCS freezing or on fear expression in RC trained mice (**Figure 3E**). In contrast, a significant impairment in fear-extinction memory was observed in EXT mice trained in the presence of *ADRAM* knockdown (**Figure 3E**), with no effect on anxiety-like behaviour in the open field test (**Supplemental Figure 7**). Together, these data demonstrate a necessary role for ADRAM in regulating Nr4a2 expression and the formation of fear-extinction memory, which is clearly the result of an effect on cognition and memory that is independent of an effect on anxiety-like behaviour.

**Figure 3.**
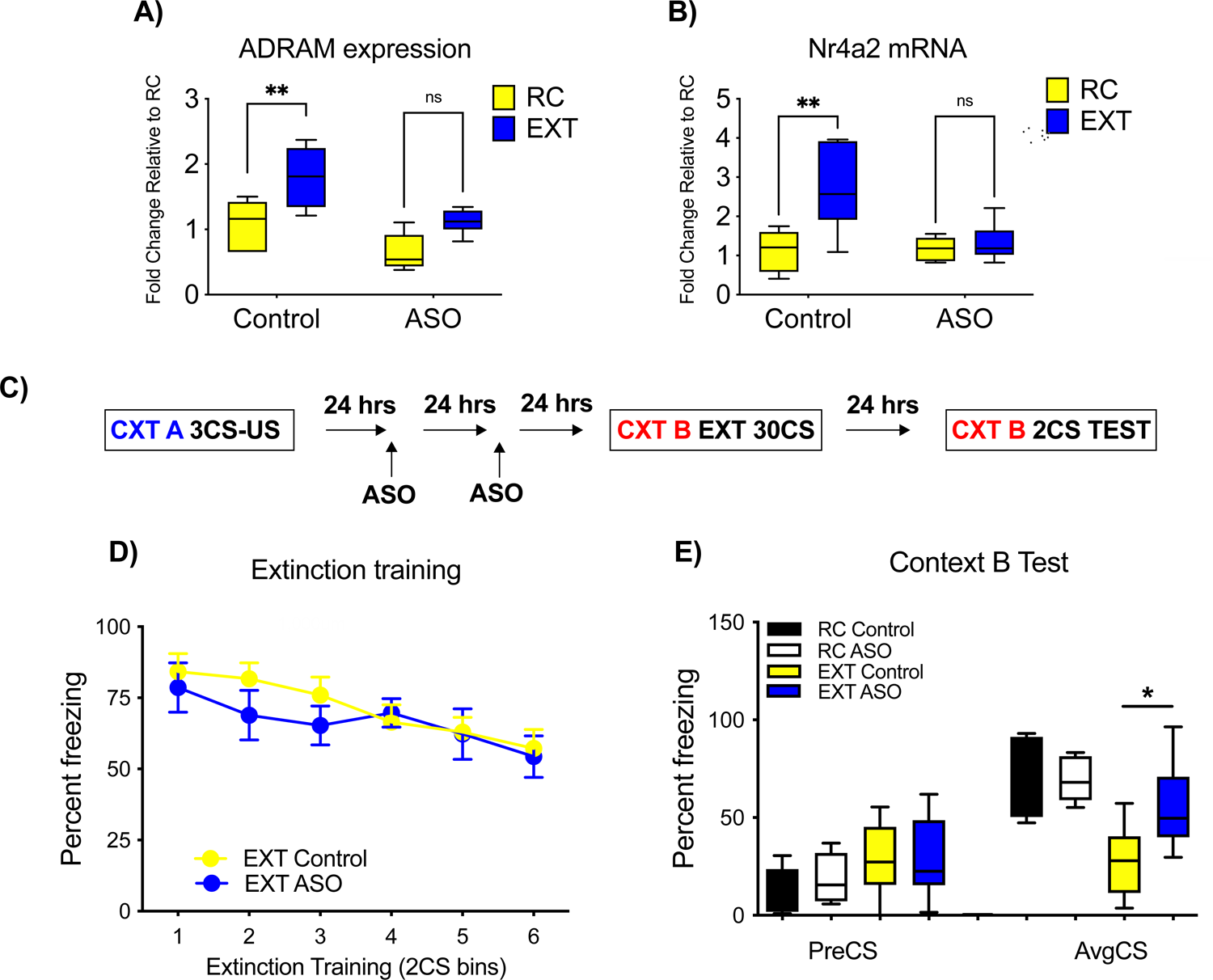
Learning-induced ADRAM expression is necessary for the induction of the immediate early gene Nr4a2 and is required for fear extinction memory. A) ADRAM ASO blocks the extinction learning-induced induction of ADRAM expression (n=5-6 biological replicates per group, two-way ANOVA, F (1, 17) = 13.18, Šídák’s post hoc analysis, Con RC vs. Con EXT, p=0.0070) and B) Nr4a2 mRNA expression (n=6 biological replicates per group, two-way ANOVA, F (1, 20) = 5.989, Šídák’s post hoc analysis, Con RC vs. Con EXT, p=0.0012) C) Schematic of the behavioral protocol used to test the effect of ADRAM ASO in the ILPFC on fear extinction memory. D) There were no significant differences between the ADRAM ASO and control groups during fear acquisition, and no effect of ADRAM ASO on preformance during within-session extinction training. E) However, knockdown of ADRAM led to a significant impairment in memory for fear extinction (n=8 animals per group, one-way ANOVA, F (3, 14) = 8.098, Šídák’s post hoc analysis, Con EXT vs. ASO EXT, p=0.0313). CS-Conditoned stimulus; PreCS- a 2 min acclimation pretest period to minimise context generalization; AvgCS-Average of 2 tone CS exposures, at test. Error bars represent S.E.M. *p<0.05, **p<0.01.

### *ADRAM* regulates the induction of the immediate early gene Nr4a2 via a direct interaction with the *Nr4a2* promoter

A substantial proportion of lncRNAs were identified as putative eRNAs based on the observation that they are transcribed from open chromatin regions of the genome and share H3K27^ac^, H3K4^me1^ and CBP features of enhancer elements. eRNAs are known to be critical for regulating adjacent protein-coding gene expression (Kim et al, 2015), although enhancer elements themselves can influence proximal gene expression through dynamic changes in the 3D architecture of the genome (Wang et al., 2019). We therefore asked if the *ADRAM* lncRNA forms a tether with the *Nr4a2* promoter through chromatin looping. Chromatin conformation capture (3C) (Hagege et al., 2007) was used to analyse the three-dimensional organization of the chromatin environment by quantifying the physical interaction between the enhancer region that encodes *ADRAM* and the *Nr4a2* gene promoter. This analysis revealed no significant change in chromatin conformation after EXT training, relative to RC (**Figure 4A, Supplemental Figure 8**), indicating that ADRAM does not coordinate the induction of Nr4a2 mRNA expression in response to fear-extinction learning via a long-distance DNA-DNA interaction.

**Figure 4.**
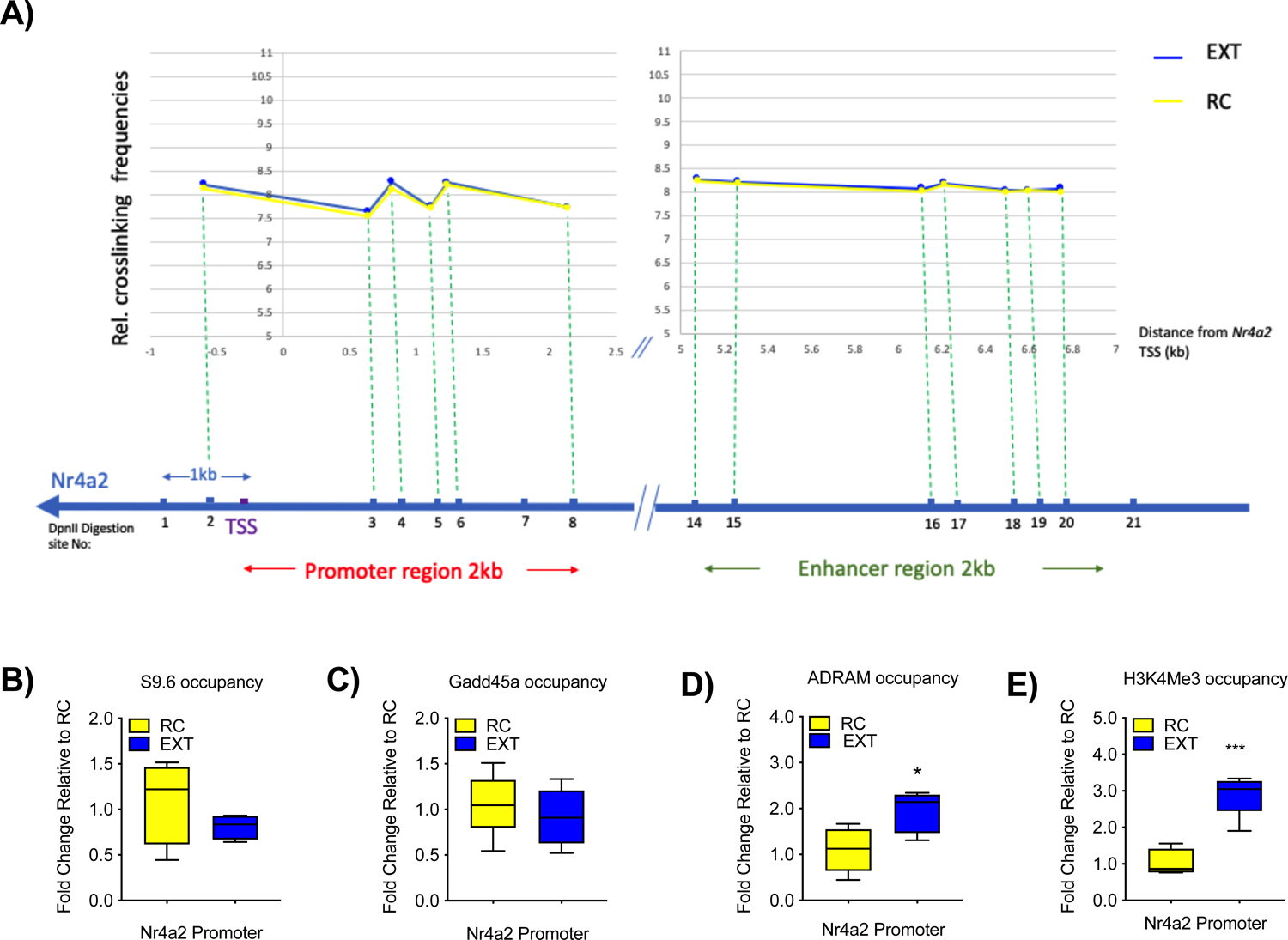
ADRAM regulates the induction of the immediate early gene Nr4a2 via a direct interaction with the Nr4a2 promoter. A) 3C-qPCR analysis of long-distance interactions at the mouse NR4A2 locus. The relative level of each ligation product (fragments −1 to 8 and 14 to 20) has been plotted according to its distance (in kb) from the Nr4a2 promoter. The data were normalized to Gapdh. Below the graphs, the TaqII restriction fragments are indicated. TaqII fragments are numbered from fragment −1 to 21. B) DRIP experiment using S9.6 antibody showing no difference in S9.6 occupancy at Nr4a2 promoter region in RC compared to EXT mice, C) There is no significant difference in Gadd45a occupancy at Nr4a2 promoter, D) ChIRP experiment demonstrating ADRAM binding to the Nr4a2 promtoer in EXT mice (n=5 biological replicates per group, two-tailed unpaired Student’s t-test), which is accompied by an increase in H3K4me3 occupancy E) (n=5 biological replicates per group, two-tailed unpaired Student’s t-test). Error bars represent S.E.M. *p<0.05, ***p<0.001.

Since eRNAs can form R-loops, which then serve to promote gene expression (Cloutier et al., 2016), we also considered the possibility that *ADRAM* forms a lncRNA:DNA hybrid, which would interact with the *Nr4a2* gene locus and regulate mRNA transcription in response to fear extinction training. To determine whether there is an R-loop structure around the transcription start site (TSS) of the *Nr4a2* promoter, we used a DNA-RNA immunoprecipitation (DRIP) assay using S9.6 antibody, which recognizes R-loops. DRIP-qPCR targeting the TSS revealed no difference between RC and EXT mice (**Figure 4B**). The effect of fear extinction learning on the occupancy of the R-loop reader protein GADD45a (Arab et al., 2019) at the same site within the *Nr4a2* promoter was also examined and, again, no evidence of R-loop formation in response to extinction training was observed (**Figure 4C**). We conclude that *ADRAM* does not form R-loops to coordinate fear extinction learning-induced *Nr4a2* mRNA expression.

Finally, to determine whether the *ADRAM* lncRNA interacts with the *Nr4a2* gene promoter, we used chromatin isolation by RNA purification (ChIRP) (Chu et al, 2015). We found that *ADRAM* binds directly to the *Nr4a2* promoter at a specific site ∼500bp upstream of the TSS (**Figure 4D, Supplemental Figure 9**), which was associated with an increase in the accumulation of the histone modification H3K4^me3^, a marker of gene activation. Critically, this effect was blocked in the presence of ASO-mediated ADRAM knockdown (**Figure 4E**). These findings strongly suggest that there is a relationship between the physical interaction between *ADRAM* and the *Nr4a2* promoter, and that this is associated with the induction of an active chromatin state.

### *ADRAM* serves as both a guide and a scaffold to coordinate fear extinction learning-induced Nr4a2 mRNA expression

LncRNAs are known to act as guides and scaffolds that recruit transcription factors and chromatin-modifying enzymes to specific gene loci. To determine whether the *ADRAM* lncRNA functions as a guide for transcriptional factors, we sought to identify the proteins that interact with *ADRAM* in response to fear extinction training. To investigate this, we performed ChIRP followed by mass spectrometry (ChIRP-MS) to provide a comprehensive profile of proteins that bind to the ADRAM lncRNA, and which could then be recruited to the *Nr4a2* promoter in response to fear-extinction learning **(Supplemental Figure 11)**. In RC mice, the most abundant interacting protein was the adhesion molecule γ-catenin, which has been shown to be a core component of the blood brain barrier (**Figure 5A, Supplemental Table 4**). Notably, the chaperone protein, *14-3-3*, was identified as the top protein to interact with the *ADRAM* lncRNA in EXT-trained nice (**Figure 5B, Supplemental Table 4**). The 14-3-3 family is highly conserved, enriched in the brain, and plays an important role in the intracellular localization of target proteins (Giles et al., 2003; Zhang et al., 2018,). We therefore validated the interaction between ADRAM and 14-3-3 by formaldehyde-RNA immunoprecipitation (fRIP) (**Figure 5C**). We then used chromatin ChIRP analysis to examine whether 14-3-3 physically interacts with the N4ra2 promoter revealing that this interaction is under temporal control. 14-3-3 interacts with the Nr4a2 promoter immediately after extinction training, but this interaction is reduced 1hr later (**Figure 5D**). Our results confirm a functional role for 14-3-3 in the brain and provide the first evidence to suggest that the 14-3-3 chaperone also acts as a temporally regulated RNA binding protein in the brain during fear-extinction learning, and that 14-3-3 is directed to the Nr4a2 promoter by ADRAM in a learning-dependent manner.

**Figure 5.**
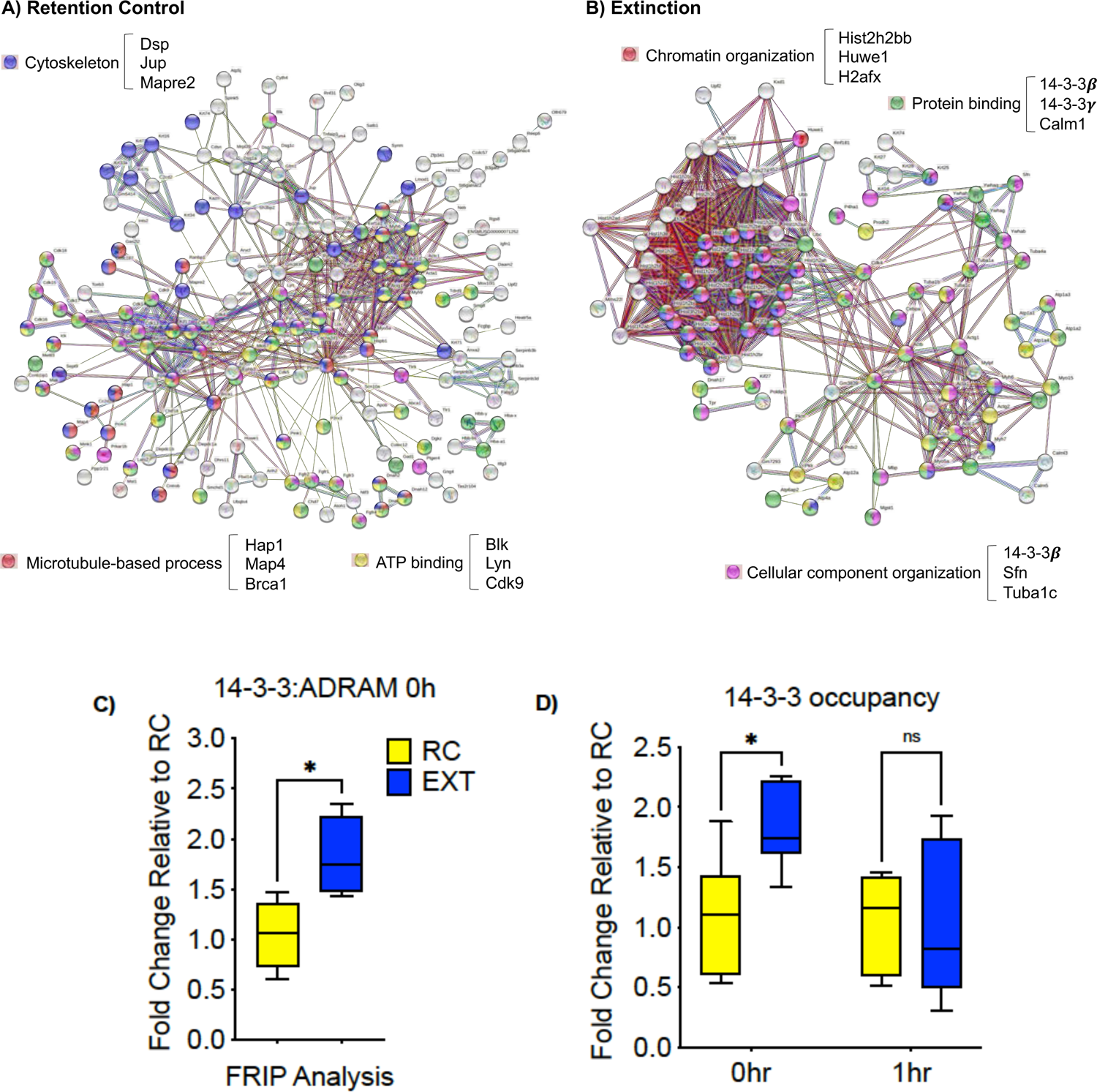
ADRAM serves as both a guide and a scaffold to coordinate fear extinction learning-induced occupancy of 14-3-3 at Nr4a2 promoter region. Representative functional interaction networks analysis of ADRAM interacting proteins in A) RC and B) EXT. C) 14-3-3β/α fRIP shows increased binding of 14-3-3β/α to ADRAM in EXT mice (n=5 biological replicates per group, two-tailed unpaired Student’s t-test). D) 14-3-3β/α chIP-qPCR reveals a transient change 14-3-3 occupancy at the Nr4a2 promoter post extinction training (n=6 biological replicates per group, two-way ANOVA, F (1, 20) = 4.589, Šídák’s post hoc analysis, 0 hr RC vs. 0 hr EXT, p=0.0309). Error bars represent SEM. * p<0.05.

We next explored the functional consequences of the activity-dependent recruitment of 14-3-3 to the Nr4a2 promoter on the epigenetic regulation of Nr4a2 gene expression. Given the critical role of HDAC3 and HDAC4 as negative regulators of learning and memory (McQuown et al., 2011; Sando et al., 2012; Zhu et al., 2019), we first questioned whether there are dynamic changes in HDAC occupancy in response to fear-extinction learning. There was a time-dependent reduction in the presence of both HDAC3 (**Figure 6A**) and HDAC4 (**Figure 6B**) at the Nr4a2 promoter following fear-extinction training. It is well established that the histone acetyltransferase CREB-binding protein (CBP) facilitates CREB activity at the *Nr4a2* promoter, which also leads to increased *Nr4a2* mRNA expression in response to various forms of learning (Bridi et al., 2017), although the mechanism by which CBP is selectively recruited has never been revealed. We found that CBP occupancy was increased at the *Nr4a2* promoter following fear extinction learning (**Figure 6C**). Following these observations, we explored whether the effect of extinction learning on HDAC3, HDAC4 and CBP occupancy at the Nr4a2 gene promoter is due, in part, to a role of ADRAM as a scaffold for 14-3-3. Indeed. Previous work has shown that the phosphorylation-dependent binding of 14-3-3 to HDAC4 serves to regulate its nuclear activity by sequestering HDAC4 from the nucleus into the cytoplasm in a signal-dependent manner (McKinsey et al., 2000; Wakeling et al, 2021).

**Figure 6.**
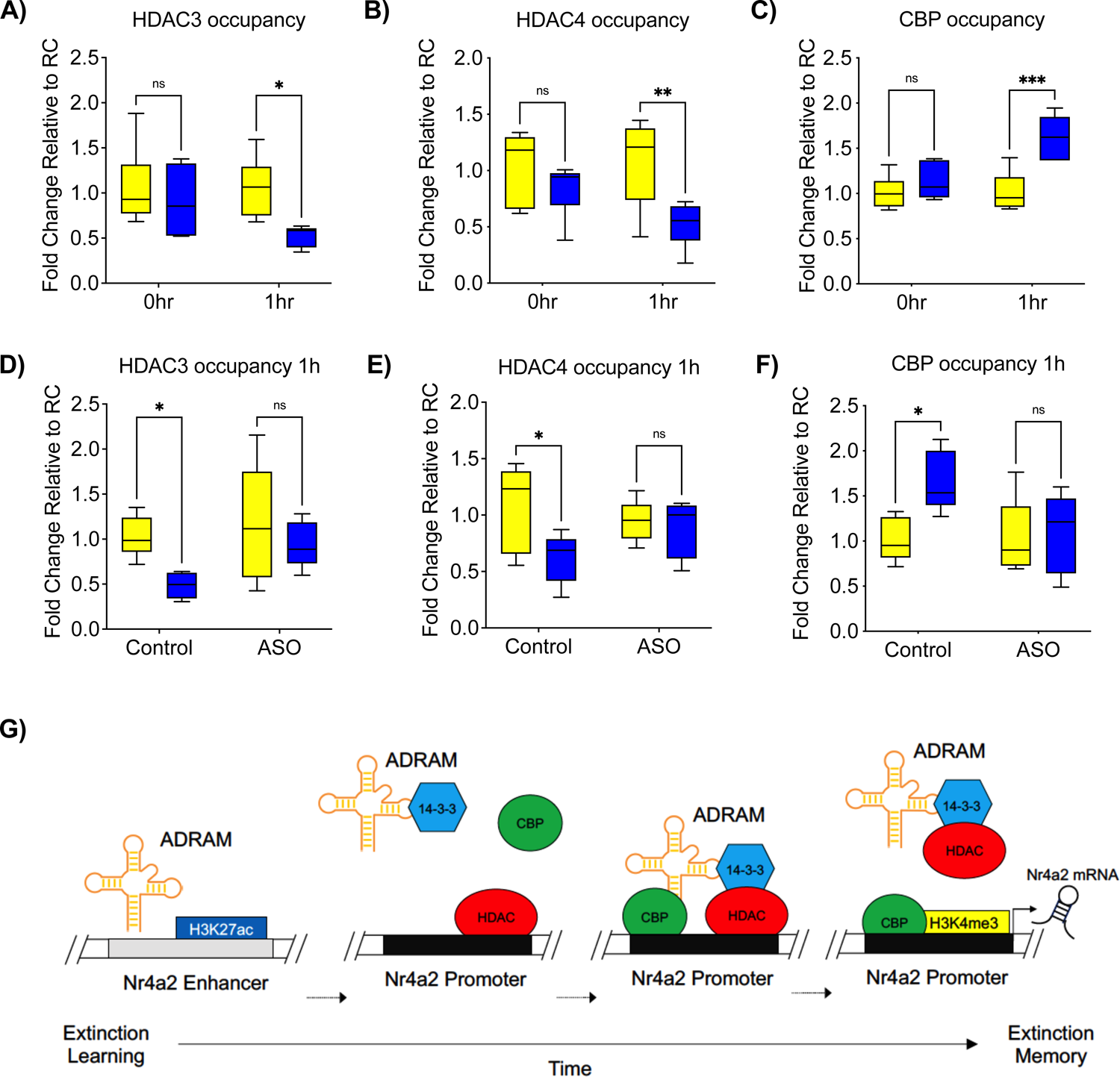
Learning-induced recruitment of 14-3-3 leads to a time-dependent change in the activity of chromatin modifiers at Nr4a2 promoter. A) ChIP-qPCR shows a reduction in both HDAC3 (n=5-6 biological replicates per group, two-way ANOVA, F (1, 19) = 5.900, Šídák’s post hoc analysis, 1 hr RC vs. 1 hr EXT, p=0.0328) and B) HDAC4 at the Nr4a2 promoter 1-hour post EXT training (n=5-6 biological replicates per group, two-way ANOVA, F (1, 19) = 10.06, Šídák’s post hoc analysis, 1 hr RC vs. 1 hr EXT, p=0.0001). C) In contrast, there was a significant increase in CBP occupancy at the Nr4a2 promoter 1hr after extinction training (n=6 biological replicates per group, two-way ANOVA, F (1, 20) = 8.273, Šídák’s post hoc analysis, 1 hr RC vs. 1 hr EXT, p=0.0328). All of the above effects were prevented in mice that had been treated with antisense oligonucleotides directed to ADRAM, D) HDAC3 (n=5-6 biological replicates per group, two-way ANOVA, F (1, 18) = 5.815, Šídák’s post hoc analysis, Con RC vs. Con EXT, p=0.0375), E) HDAC4 (n=5 biologically independent animals per group, two-way ANOVA, F (1, 16) = 4.339, Šídák’s post hoc analysis, Con RC vs. Con EXT, p=0.0404) and F) CBP. (n=5 biological replicates per group, two-way ANOVA, F (1, 16) =4.520, Šídák’s post hoc analysis, Con RC vs. Con EXT, p=0.0291). G) Proposed model of ADRAM-mediated regulation of epigenetic machinery underlying fear-extinction learning-induced Nr4a2 expression. Error bars represent S.E.M. *p<0.05.

To confirm a scaffolding role for ADRAM, an infusion of ASO into the ILPFC was used to knockdown *ADRAM* expression, which blocked the expulsion of HDAC3 and HDAC4 and inhibited the accumulation of CBP at the Nr4a2 promoter (**Figure 6D-F**). These results suggest that *ADRAM* guides 14-3-3 to the Nr4a2 promoter, which results in the time-dependent removal of HDAC3 and HDAC4, followed by the coordinated deposition of CBP. Together with the establishment of an active chromatin state in response to extinction learning, our findings confirm previous observations of a tight relationship between HDAC3, HDAC4 and CBP in the epigenetic regulation of *immediate early gene expression*, and support a new model whereby an eRNA-mediated mechanism drives the activation of *Nr4a2 in response to fear extinction learnin*g (**Figure 6G**). Since ADRAM is also expressed in other brain regions, including the hippocampus, cerebellum and somatosensory cortex (**Supplemental Figure 12**), it is likely that this eRNA is more generally involved in the regulation of experience-dependent Nr4a2 expression and in other memory processes.

### *Nr4a2* expression is necessary for the formation of fear extinction memory

Finally, given the causal effect of *ADRAM* to act as a guide and scaffold to coordinate the epigenetic regulation of extinction learning-induced *Nr4a2* mRNA expression, we wished to determine whether *Nr4a2* itself is critical for the formation of fear extinction memory. We first generated *Nr4a2* lentiviral shRNA plasmids and validated them in N2A cells, as well as verified transfection efficiency *in vivo* (**Figure 7A-C**). Infusion of the *Nr4a2* knockdown construct into the ILPFC prior to fear-extinction training (**Figure 7D**) had no effect on within-session extinction training (**Figure 7E**). There was a significant impairment in fear-extinction memory (**Figure 7F**) with no influence on preCS freezing or the ability to express fear in RC mice. Similar to the effect observed with the ADRAM knockdown, Nr4a2 shRNA treated mice exhibited normal anxiety-like behaviour (**Supplemental Figure S13**). These data that demonstrate that the effect of Nr4a2 knockdown on extinction memory is due to its influence on cognition rather than on non-specific physiological indicators of generalized anxiety. We therefore conclude that ADRAM is required for the learning-induced epigenetic regulation of Nr4a2, with our findings revealing a necessary role for the immediate early gene Nr4a2 in the formation of fear extinction memory.

**Figure 7.**
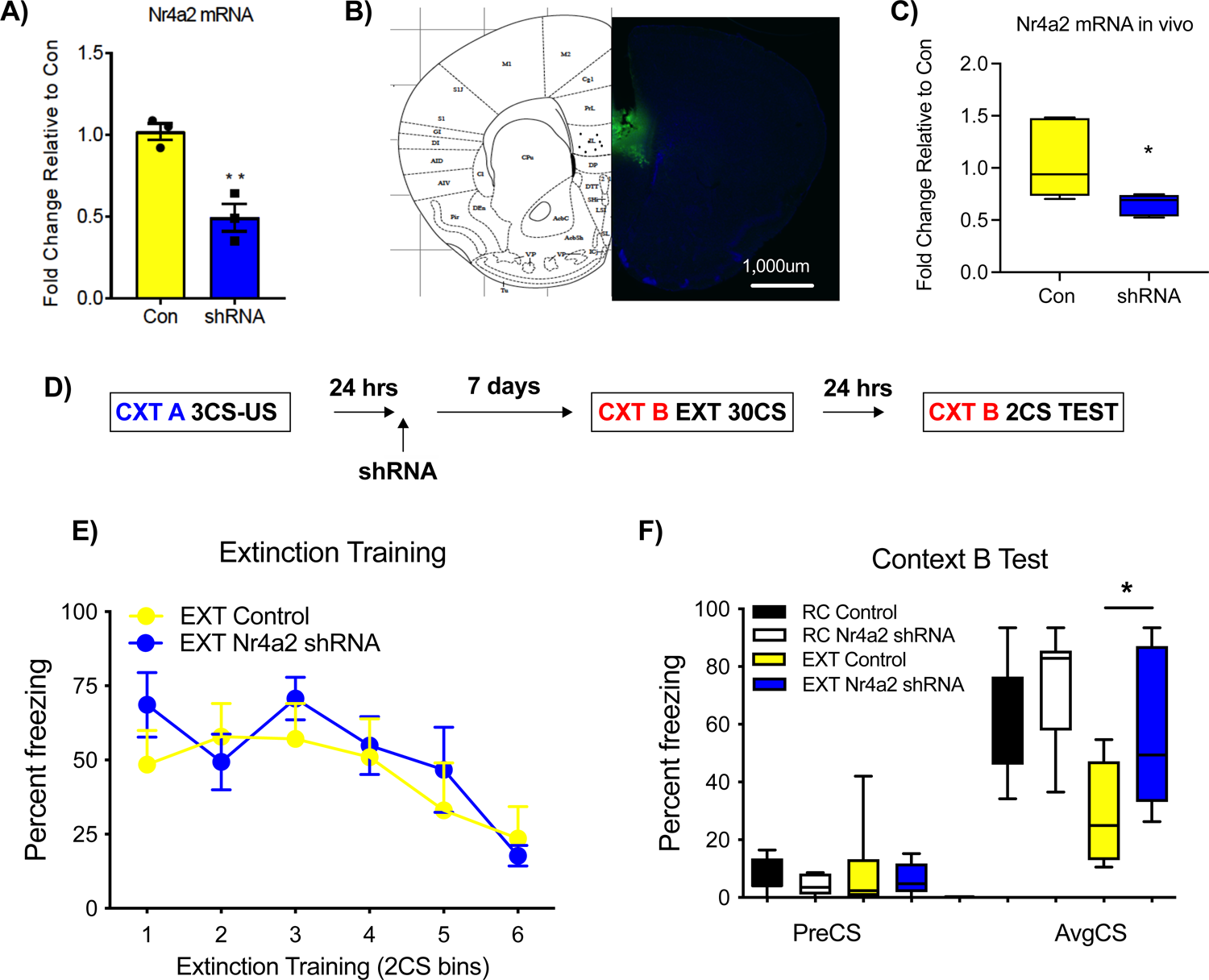
Nr4a2 expression is necessary for the formation of fear extinction memory. A) Nr4a2 shRNA decreases Nr4a2 expression in N2A cells, *in vitro* (n=3 biological replicates per group, two-tailed unpaired Student’s t-test). B) Representative image of viral infection of Nr4a2 shRNA lentivirus into the ILPFC. C) RT-qPCR shows significant Nr4a2 mRNA knockdown following Nr4a2 shRNA injection in the ILPFC (n=5-6 biological replicates per group, two-tailed unpaired Student’s t-test). D) Schematic of the behavioural protocol used to test the effect of Nr4a2 shRNA on fear extinction memory. E) There was no effect of Nr4a2 knockdown on performance during within-session extinction training. F) Nr4a2 knockdown led to a significant impairment in memory for fear extinction (n=8 animals per group, one-way ANOVA, F (3, 26) = 6.574, Šídák’s post hoc analysis, Con EXT vs. shRNA EXT, p=0.0496). CS-Conditioned stimulus; PreCS- a 2 min acclimation pretest period to minimise context generalization; AvgCS-Average of 2 tone CS exposures, at test. Error bars represent S.E.M. *p<0.05, **p<0.01.

## DISCUSSION

Here we report the discovery of widespread experience-dependent lncRNA activity in the adult ILPFC, and further reveal a significant number of inducible eRNAs that respond selectively to fear-extinction learning. This class of lncRNA was first discovered at scale more than a decade ago by the Greenberg group who identified thousands of sites outside known promoter regions in primary cortical neurons stimulated with KCl *in vitro*, which exhibited features of enhancer elements including binding of CBP and the deposition of the histone modification H3K4^me1^ (Kim et al., 2010). Transcriptional activity at these sites showed a positive correlation with downstream mRNA expression, suggesting a context-specific permissive relationship between eRNAs and their proximal mRNA partners. Malik and colleagues went on to functionally characterize neuronal enhancers and identified another histone modification, H3K27^ac^, as a key marker of their active state (Malik et al., 2014). An overlay of our lncRNA capture-seq data with learning-induced enhancer signatures in the adult brain (Halder et al, 2016), as well as our cell-type-specific ATAC-seq signatures in learning-activated Arc+ neurons revealed that there are many experience-dependent lncRNAs in the ILPFC that are endowed with features of activity-inducible eRNAs. Notably, all 10 of the validated eRNA-associated protein-coding gene candidates have been shown to be involved in plasticity, suggesting that this class of lncRNA is, in general, permissively involved in the regulation of experience-dependent gene expression.

One of the most interesting findings of our study beyond the necessary role of ADRAM in fear extinction is that ADRAM binds directly to the Nr4a2 promoter; however, in doing so it does not form an R-loop or promote chromosome looping. Trans-acting lncRNAs are known to form triplex structures on double-stranded DNA using a Hoogsteen base-pairing rule in the DNA target (Li et al., 2016; Fan et al., 2020). These structures are distinct from R-loops and could represent a novel mechanism by which lncRNAs act in a combinatorial manner to simultaneously serve as both guides and scaffolds. Indeed, examination of the 1kb upstream promoter sequence of NR4A2 revealed two sites proximal to the TSS with 25 nucleotide complementary sequences found within exon III of ADRAM. Notably, these sites overlap with G-quadruplex motifs that are predicted to enable triplex formation. These findings suggest that ADRAM functions as a guide via a direct interaction with the Nr4a2 promoter and may do so via the formation of an RNA:DNA triplex a sites of structural reactivity. Future studies will investigate whether dynamic DNA structure states are the key to how lncRNAs find their genomic targets to regulate gene expression in an experience-dependent manner.

The 14-3-3 family of evolutionarily conserved chaperone proteins is ubiquitously expressed in the brain and highly enriched at the synapse (Martin et al., 1994) being involved in a variety of neuronal processes, including synaptic plasticity (Berg et al., 2003; Marzinke et al., 2013; Foote et al., 2015). Our discovery of a direct interaction between ADRAM and 14-3-3 extends the functional capabilities of this class of chaperones to include functional activity as both an RNA binding protein and a molecule that exerts its influence through protein-protein interactions. This is not without precedent many proteins are capable of interacting with RNA, DNA, and other proteins. For example, YY1 interacts with both RNA and DNA as well as other proteins to promote its role as regulator of the Xist locus (Jeon and Lee, 2011). Finally, together with the observation that 14-3-3 is involved in learning and memory (Qiao et al., 2014), and our demonstration of how 14-3-3 interacts with eRNA to facilitate gene expression in fear extinction, these findings advance the understanding of the functional importance of this class of chaperones in the brain.

Histone modifications at neuronal enhancers also appear to be a requirement for the induction of activity-dependent genes and particularly important for rapidly induced immediate early genes (Chen et al., 2019). We found a broad overlap with H3K27^ac^, an open chromatin ATAC signature in activated neurons, and the expression of lncRNAs. Previous work has shown that eRNA activity often precedes, and then drives, the expression of the immediate early genes such as c-Fos, which occurs via a direct interaction with the histone acetyltransferase domain of CBP (Carullo et al., 2020). In addition, a large number of eRNAs have been shown to bind to CBP, correlating with the expression of downstream genes that require CBP for their induction (Bose et al., 2017). Our data on the functional relationship between ADRAM, HDAC3, HDAC4, CBP and Nr4a2 agree with these observations and, importantly, extend the findings to include the ILPFC where they are critically involved in fear extinction. Our conclusion is that ADRAM functions as both a guide and a scaffold to epigenetically regulate extinction learning-induced Nr4a2 expression. There are now many examples of multifunctional lncRNAs. For example, in dopaminergic neurons, antisense Uchl1 regulates the expression of Uchl1 in the nucleus and then shuttles to the cytoplasm where it promotes Uchl1 translation (Carrieri et al., 2012). Further, owing to its modular domain structure, Neat1 functions *in cis* to coordinate the deposition of learning-related repressive chromatin modifiers along the genome (Butler et al., 2019) and *in trans* to govern paraspeckle assembly by influencing phase separation (Yamazaki et al., 2018).

In summary, the discovery of a lncRNA that is required for fear extinction deepens our understanding of learning-induced epigenetic mechanisms by integrating the modular function of enhancer-derived lncRNAs with key epigenetic processes involved in memory, and answers the longstanding question of how certain HDACs and CBP coordinate to confer their influence on localized gene regulation with a high degree of state-dependent selectivity. LncRNAs therefore provide a bridge to link dynamic environmental signals with epigenetic mechanisms of gene regulation. Together, these findings broaden the scope of experience-dependent lncRNA activity, and underscore the importance of considering eRNAs in the adult cortex as potential therapeutic targets for fear-related neuropsychiatric disorders.

## MATERIALS AND METHODS

### Animals

10-12-week-old C57BL/6 male mice were housed four per cage, maintained on a 12hr light/dark time schedule, and allowed free access to food and water. All experiments took place during the light phase in red-light-illuminated testing rooms following protocols approved by the Institutional Animal Care and Use Committee of the University of Queensland.

### DNA/RNA Extraction

Tissue derived from the infralimbic prefrontal cortex of RC or EXT trained mice was homogenized using a Dounce tissue grinder in 500 ul of cold 1X DPBS (Gibco). 400 ul of homogenate was used for DNA extraction, and 100 ul was used for RNA extraction. DNA extraction was carried out using the DNeasy Blood & Tissue Kit (Qiagen) with RNAse A (5 prime), and RNA was extracted using Trizol reagent (Invitrogen). Both extraction protocols were followed according to the manufacturer’s instructions. The concentration of DNA and RNA was measured by Qubit assay (Invitrogen).

### Quantitative RT-PCR

Total 500 ng RNA was used for cDNA synthesis using the PrimeScript Reverse Transcription Kit (Takara). Quantitative PCR was performed on a RotorGeneQ (Qiagen) cycler with the SYBR-Green Master mix (Qiagen) by using primers for target genes and beta-actin as an internal control (**Supplemental Table 5**). The ΔΔCT method was used for analysis, and each PCR reaction was run in duplicate for each sample and repeated at least twice.

### Nr4a2 knockdown

Lentiviral plasmids were generated by inserting either Nr4a2 shRNA or scrambled control fragments (Table S1) in a modified FG12 vector (FG12H1, derived from the FG12 vector originally provided by David Baltimore, CalTech) as previously described (Li et al, 2019). Lentivirus was prepared and maintained according to protocols approved by the Institutional Biosafety Committee at the University of Queensland.

### Cannulation surgery and lentiviral infusion

A double cannula (PlasticsOne) was implanted in the anterior posterior plane, +/- 30 degrees along the midline, into the infralimbic prefrontal cortex (ILPFC). The coordinates of the injection locations were centered at +1.80 mm in the anterior-posterior plane (AP), and −2.7 mm in the dorsal-ventral plane (DV). Mice were first fear conditioned, followed by 2 ASO or lentiviral infusions (1 24 hours post fear condition training, and after a one-week incubation period, then extinction trained.

### Behavioural training and tissue collection

Two contexts (A and B) were used for all behavioral fear testing. Both conditioning chambers (Coulbourn Instruments) had two transparent walls and two stainless steel walls with a steel grid floors (3.2 mm in diameter, 8 mm centers); however, the grid floors in context B were covered by flat white plastic non-transparent surface with two white LED lights to minimize context generalization. Individual digital cameras were mounted in the ceiling of each chamber and connected via a quad processor for automated scoring of freezing measurement program (FreezeFrame). Fear conditioning was performed in context A with spray of vinegar (10% distilled vinegar). Then, actual fear condition protocol began with a 120 sec pre-fear conditioning period, followed by three pairings of a 120 sec, 80dB, 16kHZ pure tone conditioned stimulus (CS) co-terminating with a 1 sec (2 min intervals), 0.7 mA foot shock (US). Mice were counterbalanced into equivalent treatment groups based on freezing during the third training CS. For extinction (EXT), mice were exposed in context B with a stimulus light on and spray of Almond (10% Almond extracts and 10% ethanol). Mice allowed to be acclimated for 2 min, and then, extinction training comprised 30 non-reinforced 120 sec CS presentations (5-sec intervals). For the behaviour control experiments, animal only exposed into B with equal times of mice spend there by extinguished mice but were not exposed 30CS. For the retention test (RC), all mice were returned to context B and following a 2 min acclimation (used to minimize context generalization), freezing was assessed during three 120 sec CS presentations (120 sec intertrial interval). Memory was calculated as the percentage of time spent freezing during the tests.

### Open-field test

Following completion of other behavioral testing, mice were tested in open fields to check for off-target behavioral effects that could cause a change in freezing score unrelated to memory (specifically, increased generalized anxiety or reduced spontaneous locomotion). Open-field tests were conducted in a sound-attenuated room dimly illuminated with white lighting (60 ± 3 lux). Mice were placed into the center of a white plastic open field (30×30×30 cm) and recorded with an overhead camera for 20 min. Videos were analyzed using Noldus EthoVision 11 to determine the distance traveled, and the number of entries into and cumulative time spent in the center of the arena (defined as a 15×15 cm square concentric with the base of the arena).

### Behavioural Training (for tissue collection)

**Fear** conditioning consisted of three pairing (120 sec inner-trial interval ITI) of a 120 sec, 80dB, 16kHZ pure tone conditioned stimulus (CS) Co-terminating with a 1 sec, 0.7 mA foot shock in context A. Mice were matched into equivalent treatment groups based on freezing during the third training CS. Context A exposure group spent an equivalent amount of time in context A without any CS and US. One day later, the fear-conditioned mice were brought to context B, where the extinction group (EXT) was presented with 60 CS presentations (5s ITI). The retention control (RC) group spent an equivalent amount of time in context B without any CS presentations. ILPFC was collected from both of these groups immediately after the end of either context B exposure (RC) or extinction training (EXT).

### Immunohistochemistry

Mice were euthanized with 100 mg/Kg ketamine mixed with 10 mg/Kg xylazine, after which they were perfused with 60 ml 1X PBS followed by 4% paraformaldehyde in 1X PBS. Following extraction, the brains were stored 4% paraformaldehyde overnight. The brains were then placed in 30% sucrose for a minimum 24hr prior to cryostat slicing. Sectioning at 40um was performed using CM1950 cryostat (Leica), and sections were mounted on Superfrost Plus microscope slides (Fisher Scientific). The sections were incubated 1-2hr in blocking buffer, after which primary antibodies (MAP2 or GFP) were added and the slides incubated at 4 °C overnight. The slides were then washed 3 times with 1X PBS containing 0.02 % Tween 20 (PBS-T), after which secondary antibodies were added (Dylight 488-conjugated AffiniPure sheep anti-goat IgG or Dylight 549-conjugated AffiniPure goat anti-rabbit IgG, Jackson ImmunoReasearch Laboratories). The slides were incubated at room temperature for 45 min, washed 3 times with 1X PBST and sealed with Prolong Diamond Antifade Mountant (Life technology).

### Fluorescence In Situ Hybridization

Custom Stellaris® FISH Probes were designed against lncRNA ADRAM by utilizing the Stellaris® FISH Probe Designer (Biosearch Technologies, Inc., Petaluma, CA) available online at www.biosearchtech.com/stellarisdesigner (**Supplemental Table 4**). The cultured primary neurons were hybridized with the ADRAM Stellaris FISH Probe set labeled with TAMRA (Biosearch Technologies, Inc.), following the manufacturer’s instructions.

### Chromatin immunoprecipitation

Chromatin immunoprecipitation (ChIP) was performed following modification of the Invitrogen ChIP kit protocol. ILPFC samples were fixed in 1% formaldehyde and cross-linked cell lysates were sheared by Covaris in 1% SDS lysis buffer to generate chromatin fragments with an average length of 300bp by using Peak Power: 75, Duty Factor: 2, Cycle/Burst: 200, Duration: 900 Secs and temperature: between 5 °C to 9 °C. The chromatin was then immunoprecipitated using the specific antibody to each target. Also, equivalent amount of control normal rabbit IgG (Santa Cruz) was used for non-specificity control and the sample was incubated overnight at 4°C. Protein-DNA-antibody complexes were precipitated with protein G-magnetic beads (Invitrogen) for 1hr at 4°C, followed by three washes in low salt buffer, and then three washes in high salt buffer. The precipitated protein-DNA complexes were eluted from the antibody with 1% SDS and 0.1 M NaHCO3, then incubated 4hr at 60 °C in 200 mM NaCl to reverse formaldehyde cross-link. Following proteinase K digestion, phenol-chloroform extraction, and ethanol precipitation, samples were subjected to qPCR using primers specific for 200 bp segments within corresponding target regions.

### Primary cortical neuron, N2A and HEK cell culture

Cortical tissue was isolated from E15 mouse embryos in a sterile 50ml tubes. To dissociate the tissue, it was finely chopped followed by gentle pipetting to create a single cell suspension. To prevent clumping of cells, the homogenate was treated with 2 unit/ul of DNase I. The dissociated cells were passed through a 40um cell strainer (BD Falcon) and plated onto 6 well plates coated with Poly-L-Ornithine (Sigma P2533) at a density of 1×106 cells per well. The medium used was Neurobasal media (Gibco) containing B27 supplement (Gibco). 1X Glutamax (Gibco), and 1% Pen/Strep (Sigma). N2a cell was maintained in medium containing half DMEM, high glucose (GIBCO), half OptiMEM 1 (Gibco) with 5% serum and 1% Pen/Strep. HEK293t cell was maintained in medium contains DMEM, high glucose (Gibco) with 5% serum and 1% Pen/Strep.

### ASO knockdown ADRAM in vitro

200nM of Adram ASO or scramble control (**Supplemental Table 5**) was dropped on to primary cortical neurons in a 6-well plate. After 7 days incubation, cells were harvested for RNA extraction.

### Fluorescence Activated Cell Sorting (FACS)

Frozen ILPFC tissue samples were homogenized and fixed with 1 % methanol free PFA (Thermo) at room temperature for 5 mins. Glycine, at a final concentration of 0.125 mM was used to stop the fixation reaction. The cells were then washed three times with 1X cold PBS. The cell suspension was treated with DNaseI (Thermo) for 15 mins at 4 °C followed by a wash with 1ml of 1X cold PBS. The cell suspension was then blocked using FACS blocking buffer (1X BSA, 1X normal Goat Serum and 1% TritonX) for 15 min at 4 °C with end-to-end rotation. After 15 min blocking, the cell suspension was incubated with 1:150 dilution of preconjugated Arc antibody (Bioss) and 1:300 dilution of preconjugated NeuN antibody (Bioss) at 4 °C for 1hr with an end-to-end rotation. At the end of incubation, two rounds of 1ml 1X cold PBS washes were applied. The cell pellets were resuspended with 500ul of 1X cold PBS, and a 1:2000 dilution of DAPI was added. FACS was performed on a BD FACSAriaII (BD Science) sorter within the Flow Cytometry Facility at the Queensland Brain Institute, University of Queensland, Brisbane.

### Long noncoding RNA capture sequencing

ILPFC tissue was collected immediately after RC or EXT training (N=24). 2Dg of total RNA isolated from ILPFC tissue (4 pooled per library, 3 libraries per group) was used to generate the RNA library. Total RNA was treated with NEBNext® rRNA Depletion Kit (NEB) to remove rRNA. RNA libraries were then generated by NEBNext® Ultra™ II RNA Library Prep Kit for Illumina® (NEB). A custom-designed probe panel (Roche), which targets 28,228 known and predicted mouse lncRNAs (Bussotti et al, 2016), was used to capture lncRNAs in accordance with the SeqCap EZ Hybridization and Wash Kit and SeqCap EZ Accessory Kit (Roche). Captured lncRNA libraries were sequenced on an Illumina HiSeq 4000 platform with a read length of 150 bp x 2.

### Assay for Transposase-Accessible Chromatin using sequencing (ATAC-seq)

Male mice (N=30) were either RC (n=15) of EXT (n=15) trained and five ILPFCs were pooled together for FACS. After FACS, Arc+NeuN+ and Arc-NeuN+ populations from RC or EXT group (n=3 libraries per pooled group) were used for ATAC-seq in following procedure. 50,000 cells were resuspended in 50 μl 2x lysis buffer (20 mM Tris–Cl, pH 7.4; 20 mM NaCl; 6 mM MgCl2; 0.10% Igepal CA-630) and spun down immediately (500 g for 10 min at 4 °C). The transposase reaction was performed using the ATAC-seq kit (Active motif) following the manufacturers protocol. Paired-end (PE) libraries were sequenced using an Illumina HiSeq 4000 sequencing platform with the read length of 150 bp x 2.

### Sequencing data analysis

Cutadapt (v1.17, https://cutadapt.readthedocs.io/en/stable/) was used to trim off low-quality nucleotides (Phred quality lower than 20) and Illumina adaptor sequences at the 3’ end of each read for both lncRNA capture sequencing (Capture-Seq) and ATAC sequencing (ATAC-Seq) data. Processed reads were aligned to the reference genome of mouse (mm10) using HISAT2 (v2.1.0) (Kim et al, 2015) and BWA-MEM (0.7.17) (Li and Durbin, 2009) for Capture-Seq and ATAC-Seq data, respectively. SAMtools (version 1.8) (Li et al, 2009) was then used to convert “SAM” files to “BAM” files, remove duplicate reads, sort and index the “BAM” files. To avoid the artefact signals potentially introduced by misalignments, we only kept properly PE aligned reads with mapping quality at least 20 for downstream analyses. For Capture-Seq data, three rounds of StringTie (v2.1.4) (Pertea et al, 2015) was applied to i) perform reference-guided transcriptome assembly by supplying the GENCODE annotation file (V25) with the “-G” option for each sample, ii) generated a non-redundant set of transcripts using the StringTie merge mode, and iii) quantitate the transcript-level expression for each sample, with the option of “-e -G merged.gtf”. For the lncRNA analysis, known protein-coding transcripts (with the GENCODE transcript biotype as “protein-coding”) or transcripts with a length of less than 200nt were removed from the StringTie results. Ballgown (v2.22.0) (Frazee et al, 2015) was then used to conduct transcript-level differential expression analysis. For ATAC-Seq data, BEDtools (v2.27.1) (Quinlan et al, 2010) was used to convert the BAM files to BED files. Reads aligned to the mitochondrial genome were discarded as the mitochondrial genome is more accessible due to the lack of chromatin packaging. To account for the 9-bp duplication created by DNA repair of the nick by Tn5 transposase, reads were shifted + 4 bp and − 5 bp for positive and negative strand, respectively. MACS2 (v2.2.4) (Zhang et al, 2008) was used for peak calling with the option of “--shift −75 --extsize 150 --nomodel -B --SPMR -g mm --keep-dup all”. One Arc+ EXT sample was removed from the analysis due to extremely low sequencing quality. BEDtools with the “multiIntersectBed” function was used to identify the consistent peaks among biological replicates. Only consistent ATAC peaks detected in all biological replicates in at least one condition were collected and used for downstream analyses. We categorized ATAC peaks into different genomic categories using a custom PERL script. H3K27ac, H3K4me1 and CBP data were downloaded from the NCBI GSE with the accession IDs “GSM1939159”, “GSM1939160”, “GSM1939127”, “GSM1939128” and “GSM530174”. For the CBP peaks, the genomic coordinates were converted from mm9 to mm10 using the liftOver script. LncRNAs containing each of the ATAC, H3K27ac, H3K4me1 and CBP signatures were identified using a custom PERL script.

### Code and data availability

The data analysis pipeline and customized PERL scripts are available through the GitHub page (https://github.com/Qiongyi/lncRNA_2020). All the sequencing data are publicly available with the accession IDs XXXXXX. In addition, we have also made our custom tracks publicly available via the UCSC genome browser with the link (http://genome.ucsc.edu/cgi-bin/hgTracks?db=mm10&hubUrl=https://data.cyverse.org/dav-anon/iplant/home/qiongyi/lncRNA2020/hub_lncRNA2020_v1.0.txt).

### Quantitative Analysis of Chromosome Conformation Capture Assay (3C-qPCR)

The 3C-qPCR protocol was adapted from the previous publications (Fullwood et al, 2009). Briefly, ILPFC samples were harvested from the either extinction training or control group mice and dissociated with 1x cold PBS in the douncer to generate the cell suspension. Approximal 1.5 x 106 cells from each mPFC were fixed with 1% formaldehyde for 10 min at room temperature and lysed, and then nuclei were digested with DpnII (NEB) before ligation. All primers were designed within 150 bp from DpnII digestion site (Supplemental Table 4). The specificity and amplification efficiency of each primer were tested by performing qPCR on a serial dilution of the BAC clone (cat No., which contains the Nr4a2 locus) and then generate the standard curve. Digestion efficiency, ligation efficiency, and sample purity were all verified as per established protocols (8). According to a (slope) and b (intercept) based on the standard curve of the BAC clone, we then transformed the values as 10^(Ct-b)/a for each primer and normalized to GAPDH. The 3C quantitative results are presented as the mean ± S.E.M from three independent preparations of 3C sample with duplicate qPCR data.

### Chromatin isolation by RNA purification (ChIRP) analysis

ChIRP analysis was performed as previously described (9). Briefly, 1 ml of PBP (1xPBS and 1xPIC) was added into each mouse’s ILPFC and gently homogenized. The final concentration of 3% formaldehyde solution was used to fix for 30 minutes, with glycine termination and centrifuge to get rid of supernatant. The pellet was resuspended with lysate buffer and incubate for 15mins on ices, then sheared using Covaris M220. The oligonucleotide probe mix labeled with a biotin marker for ADRAM was added to the product and hybridized at 37 degree for 4 hours. Following that, Streptomycin biotin C1 protein beads were added and incubated for half an hour, then washed for 3 times and the magnetic beads were re-suspended in the biotin elution buffer (12.5mM biotin, 7.5mM HEPES [pH7.5], 75mM NaCl,1.5mM EDTA, 0.15% SDS, 0.075% sarkosyl, and 0.02% Na-Deoxycholate). After shaking at room temperature for 20 minutes and 65 degrees for 10 minutes, put the tube on the magnetic rack. Take the supernatant for DNA extraction and qPCR using the primers amplifying the Nr4a2 promoter region.

### Chromatin purification by RNA precipitation followed by mass spectrometry (ChIRP-MS)

Similar to ChIRP, after elution with the biotin elution buffer, the TCA precipitation method was used to extract protein for mass spectrometry to identify proteins associated with ADRAM.

### HPLC/MS MS/MS analysis

Following the sample preparation method of Xiong et al (2021), peptide extracts were analysed by nanoHPLC/MS MS/MS on an Eksigent ekspert nanoLC 400 system (SCIEX) coupled to a Triple TOF 6600 mass spectrometer (SCIEX) equipped with PicoView nanoflow (New Objective) ion source. Full scan TOFMS data was acquired over the mass range 350–1800 and for product ion ms/ms 100–1500. Ions observed in the TOF-MS scan exceeding a threshold of 200 counts and a charge state of +two to +five were set to trigger the acquisition of product ion, ms/ms spectra of the resultant 30 most intense ions.

### MS data analysis

Data was acquired and processed using Analyst TF 1.7 software (SCIEX). Protein identification was carried out using ProteinPilot software v5.0 (SCIEX) with Paragon database search algorithm. MS/MS spectra were searched against the mouse proteome in the UniProt database (55,366 proteins). The search parameter was set to through with False Discovery Rate (FDR) analysis. A non-linear fitting method was used to determine both a global and a local FDR from the decoy database search (Tang et al., 2008). The cut-off for identified proteins was set to 1% global FDR. The MS2Count was calculated for each identified protein by summing the MS2Count of all peptides belonging to that protein. The proteins identified using control probe were subtracted from the list of ADARAM bound proteins and then all proteins were analyzed by STRING to determine functional protein association networks (https://string-db.org/) (**Supplemental Table 4**).

## Statistical Analyses

All statistical analysis was performed using Prism 8. Following an analysis of descriptive statistics, two-tailed unpaired Student’s t-test was used for direct comparison between RC and EXT groups at each time point. One-way or two-way ANOVA was chosen for multiple comparisons where appropriate. All multiple post hoc analysis was performed using Šídák’s multiple comparison test. Error bars represent SEM. Significant differences were accepted at p<0.05.

## Acknowledgements

The authors gratefully acknowledge grant support from the Brain and Behavioral Research Foundation (NARSAD Independent Investigator Award, TWB), NIH 1R21MH103812 (TWB), ARC DP180102998 (TWB), ARC DP190100234 (TWB), and NSFC 82001421 (XL). LJL, ELZ and SUM are supported by the Westpac Future Scholars and the University of Queensland. Imaging/Analysis was performed at the Queensland Brain Institute’s Advanced Microscopy Facility. We thank Mr. Alun Jones from the IMB Mass Spectrometry Facility for his technical support, and Ms. Rowan Tweedale for helpful editing of the manuscript.

## AUTHOR CONTRIBUTIONS

Conceptualization, T.W.B., W.W., and X.L.; Experiments, X.L., Z.Q.W., W.S.L., P.R.M., G.S., E.L.Z., L.J.L., S.U.M., A.K., H.B.R., S.J.C., and J.J.Z.; Computational analyses, D.B., and Q.Y.Z.; Data interpretation, T.W.B., W.W., Q.Y.Z., and X.L.; Writing – original draft, T.W.B., X.L. and W.W.; Writing – review & editing, T.W.B., W.W., Q.Y.Z., T.M., J.S.M. and X.L.

---

**Supplemental Figure 1.**
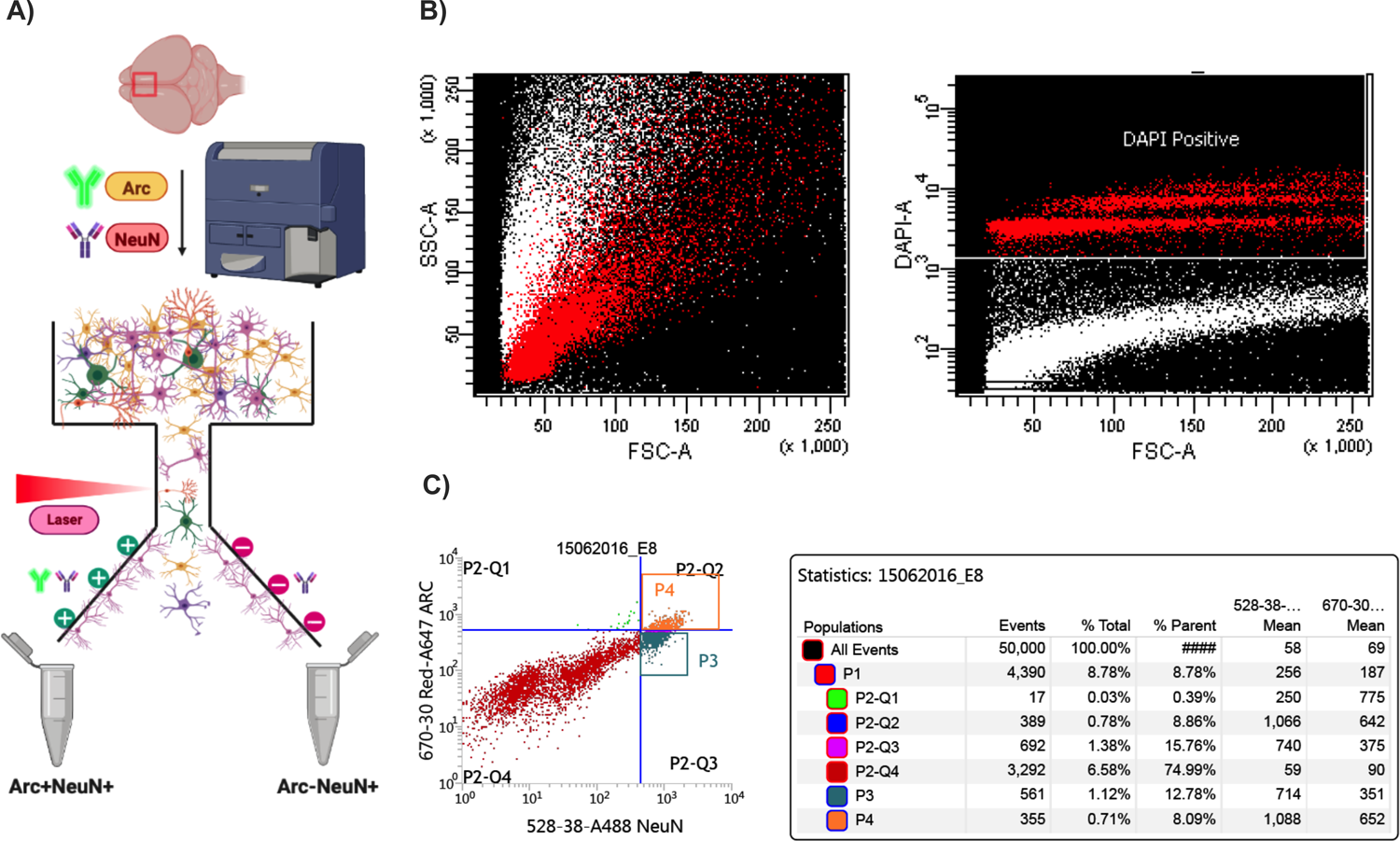
Fluorescence-activated cell sorting (FACS) to enrich for ILPFC neurons activated by leaning. A) Flow chart of experimental procedure. B) Representative images of density plots for selecting single neurons (Left: based on size selection; Right: based on DAPI staining). C) Isolating activated neurons using co-staining of Arc antibody conjugated with Alexa Fluor 647, and NeuN antibody conjugated with Alexa Fluor 488. Representative Density plot shows the FACS selection gate setting and representative statistics table shows the percentage of cells isolated based on the fluorescence signal under different selective criteria.

**Supplemental Figure 2.**
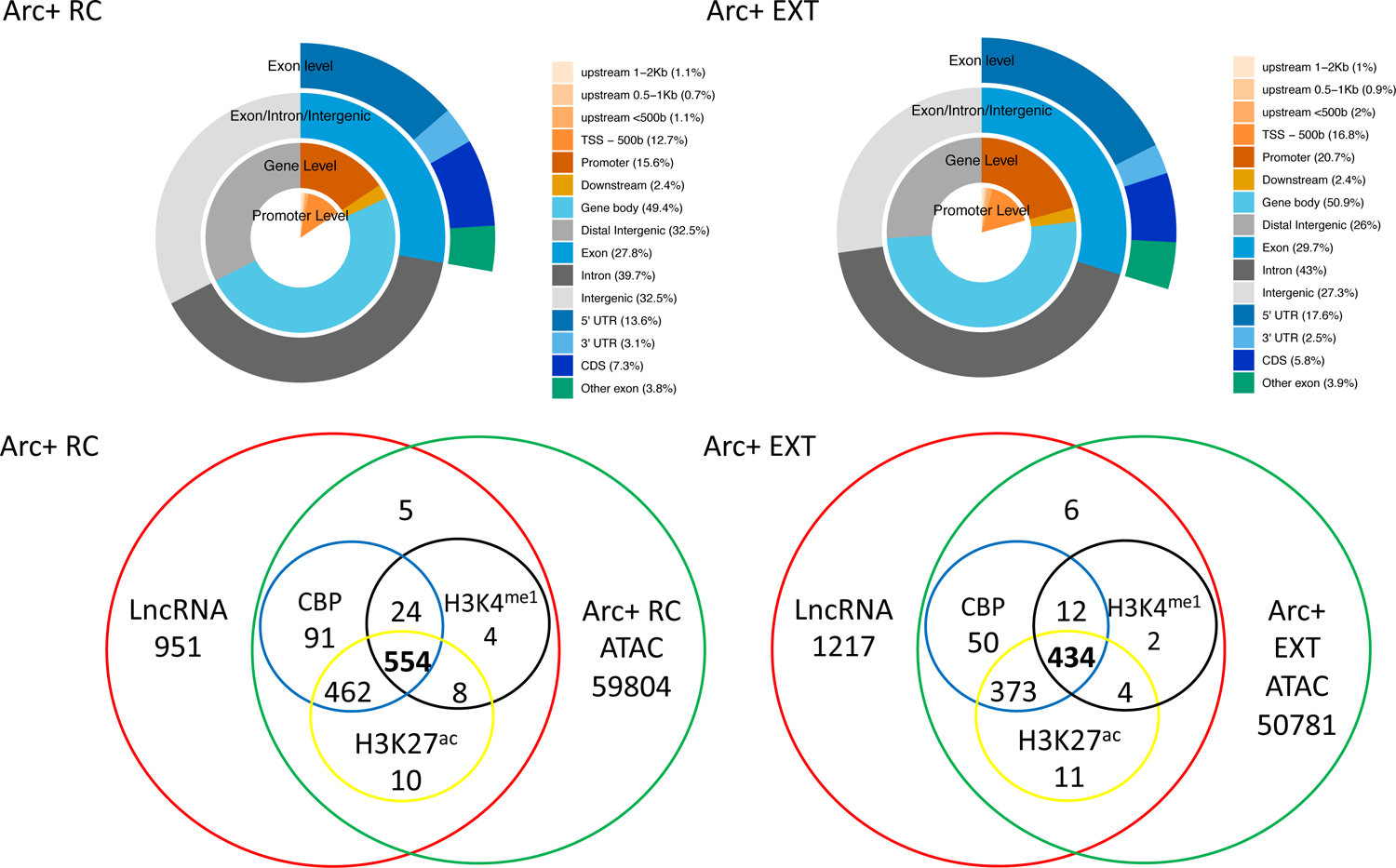
Genome wide sequencing data analysis reveals cell-type specific extinction-learning-induced eRNAs. Genomic distribution of ATAC peaks in activated neurons derived from A) retention control (RC) and B) fear extinction (EXT) trained animals. C) and D) Venn diagram showing overlap between proximal lncRNAs, ATAC-seq, H3K27^ac^, CBP and H3K4^me1^ in RC and EXT trained mice.

**Supplemental Figure 3.**
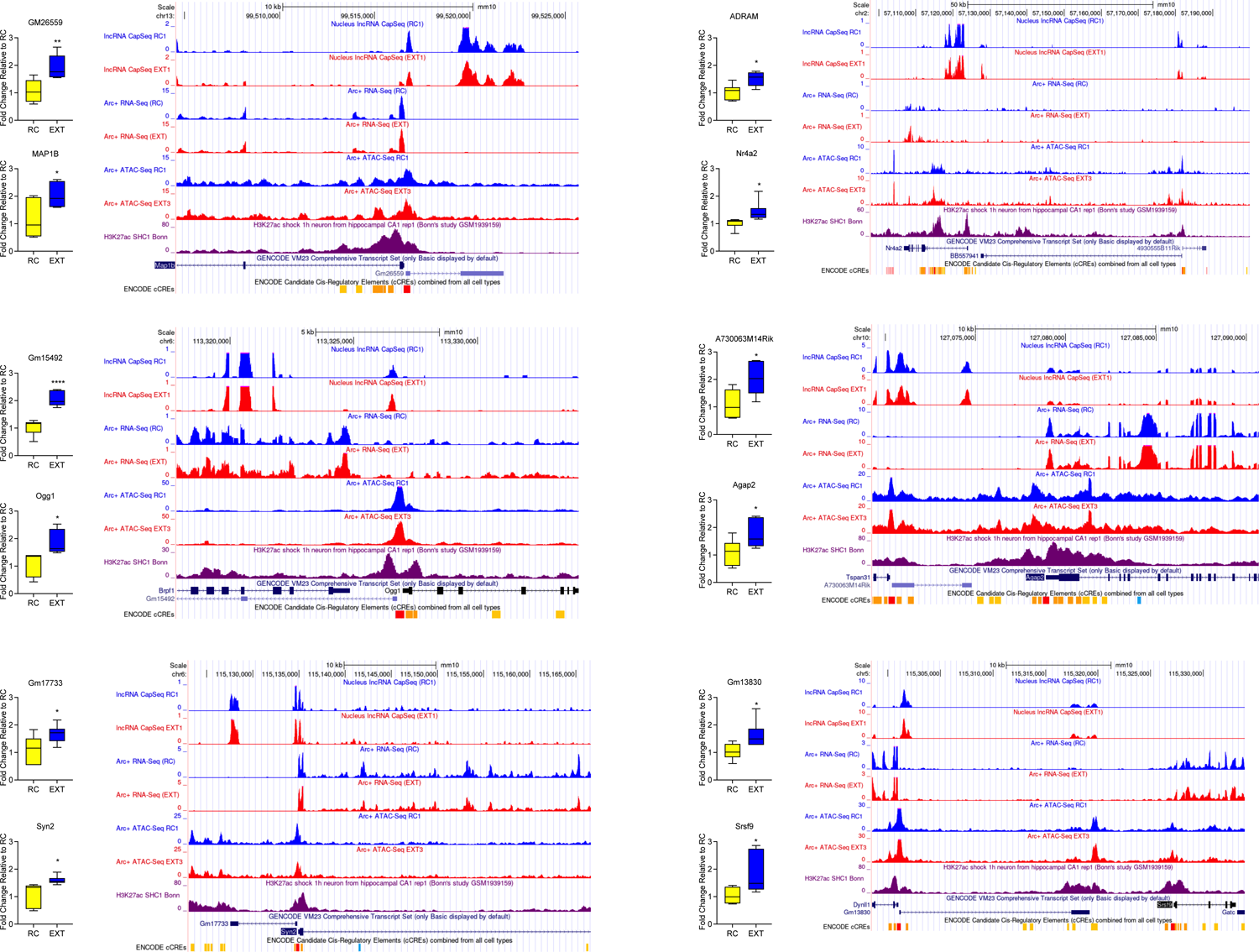
Fear-extinction learning induces eRNA and proximal protein-coding mRNA expression. Part 1: A) A730063M14Rik:Agap2, B) Gm17733:Syn2, C) Gm15492:Ogg1, D) Gm26559:Map1b, E) Gm13830:Srsf9, and F) BB557941:Nr4a2. Part 2: G) AC124502:Npas4, H) 2410080102Rik:Klf9, I) B930059L03Rik:Dync1h1 and J) GM38285:Tshz3. UCSC genome browser tracks represent the expression profile of each eRNA:mRNA pair and their overlap with publicly available genome-wide H3K27ac chIP-seq data. (n=6 independent biological replicates per group, two-tailed unpaired Student’s t-test). Error bars represent S.E.M. * p<0.05, ** p<0.01, **** p<0.0001.

**Supplemental Figure 4.**
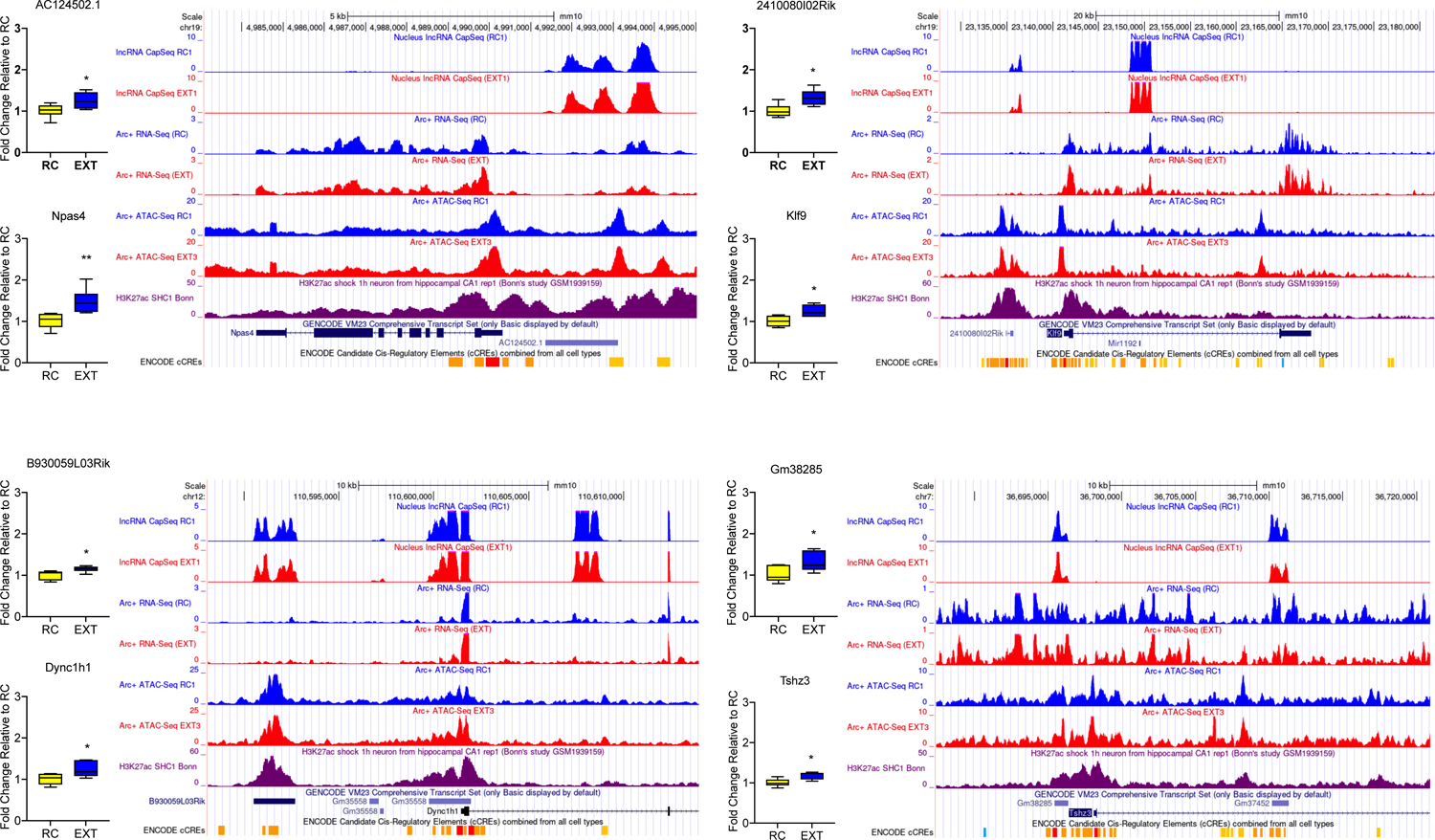
RNA-FISH image showing the subcellular localization of ADRAM in primary cortical neurons. Scale bar: 20 μm.

**Supplemental Figure 5.**
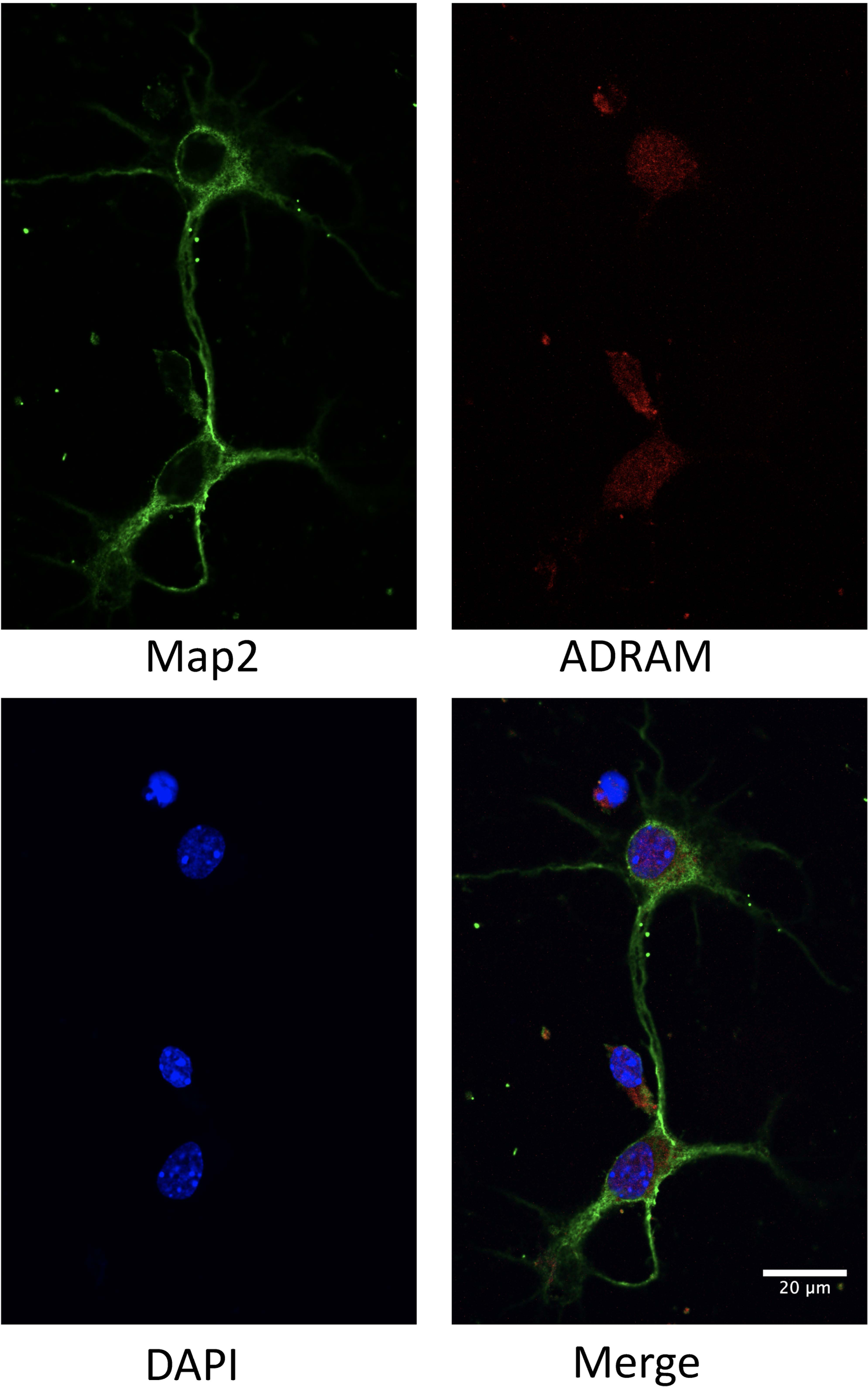
ADRAM ASO validation in primary cortical neurons. A) ADRAM ASO1, 2 and 3 lead to a reduction in ADRAM expression (n=3-4 biological replicates per group, one-way ANOVA, F (3, 11) = 12.60, Šídák’s post hoc analysis), with ASO1 showing the greatest effect. B) In primary cortical neurons, ADRAM ASO1 inhibits the expression of Nr4a2 (n=4 biological replicates per group, one-way ANOVA, F (3, 12) = 35.63, Šídák’s post hoc analysis). Error bars represent SEM. * p<0.05, ** p<0.01, *** p<0.001.

**Supplemental Figure 6.**
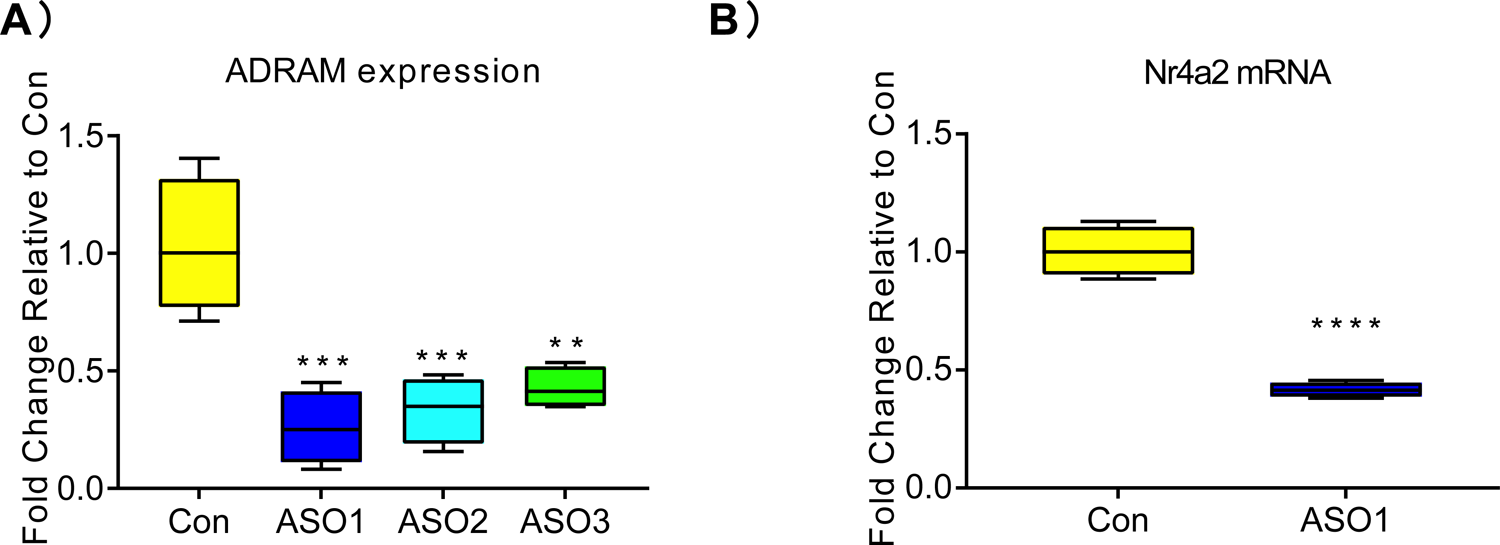
ADRAM ASO test in ILPFC. A) Representative image of ASO injection at ILPFC. B) RT-qPCR result shows effective knockdown of ADRAM by direct infusion of ASO into the ILPFC (n=5 animals per group, one-way ANOVA, F (3, 12) = 35.63, Šídák’s post hoc analysis). Error bars represent SEM. * p<0.05, ** p<0.01, *** p<0.001.

**Supplemental Figure 7.**
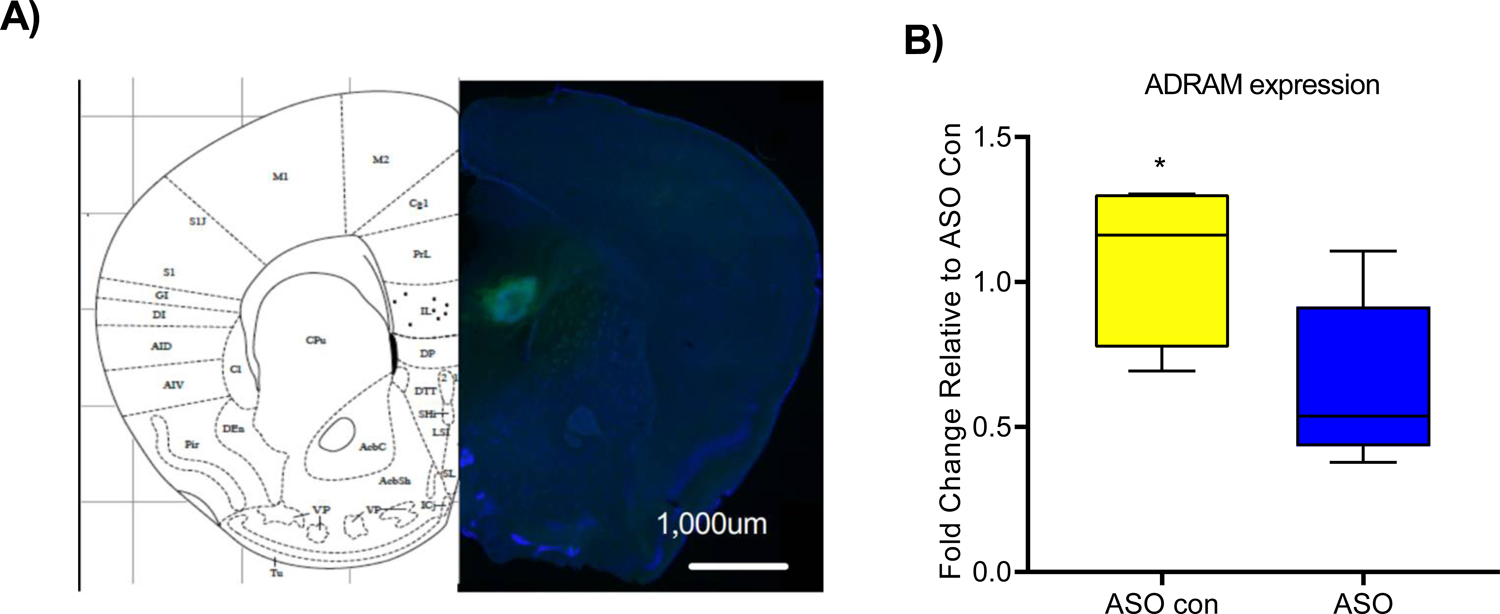
ADRAM ASO has no effect on the anxiety-like behaviour. Compared to controls, mice injected with ADRAM ASO show no differences in time spent in center A), distance travelled B) and entries to center C).

**Supplemental Figure 8.**
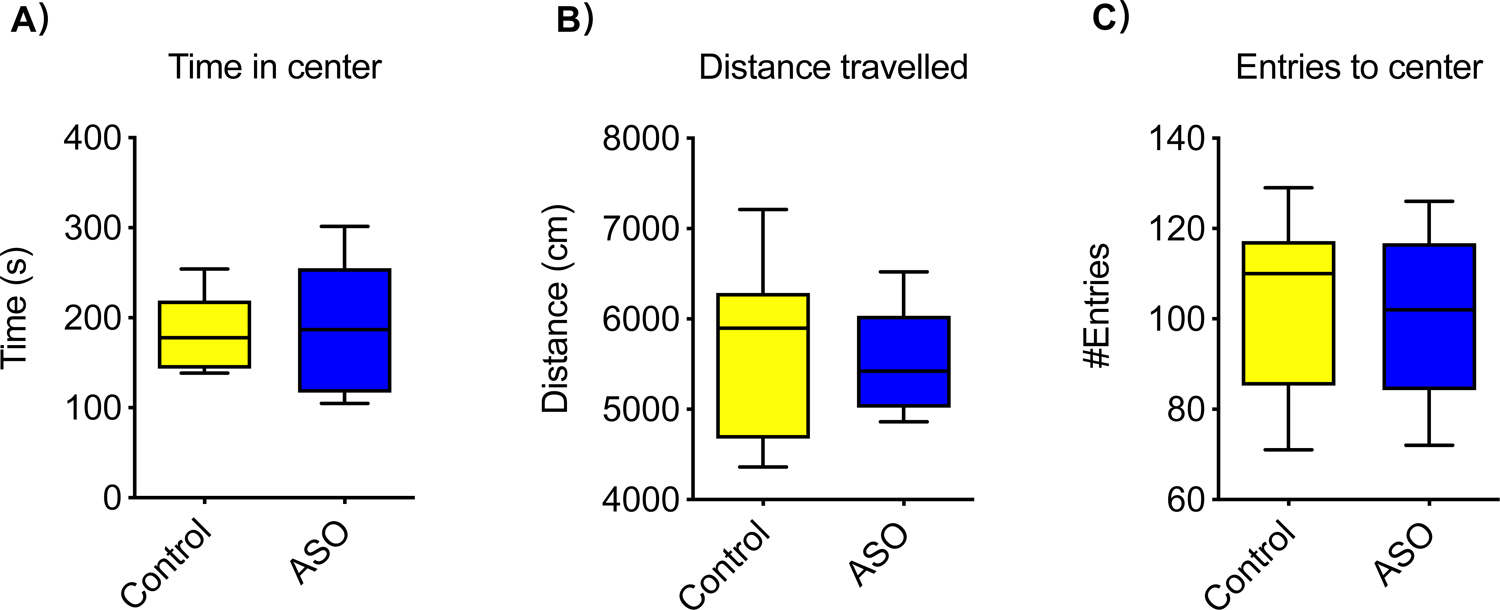
Restriction enzyme selection for 3C-qPCR experiment. A) Top, 4-base cutters Acil, DpnII and TaqI were chosed to test. Bottom) DpnII shows the best digestion efficiency and cuts 23 sites around the Nr4a2 promoter and enhancer regions.

**Supplemental Figure 9.**
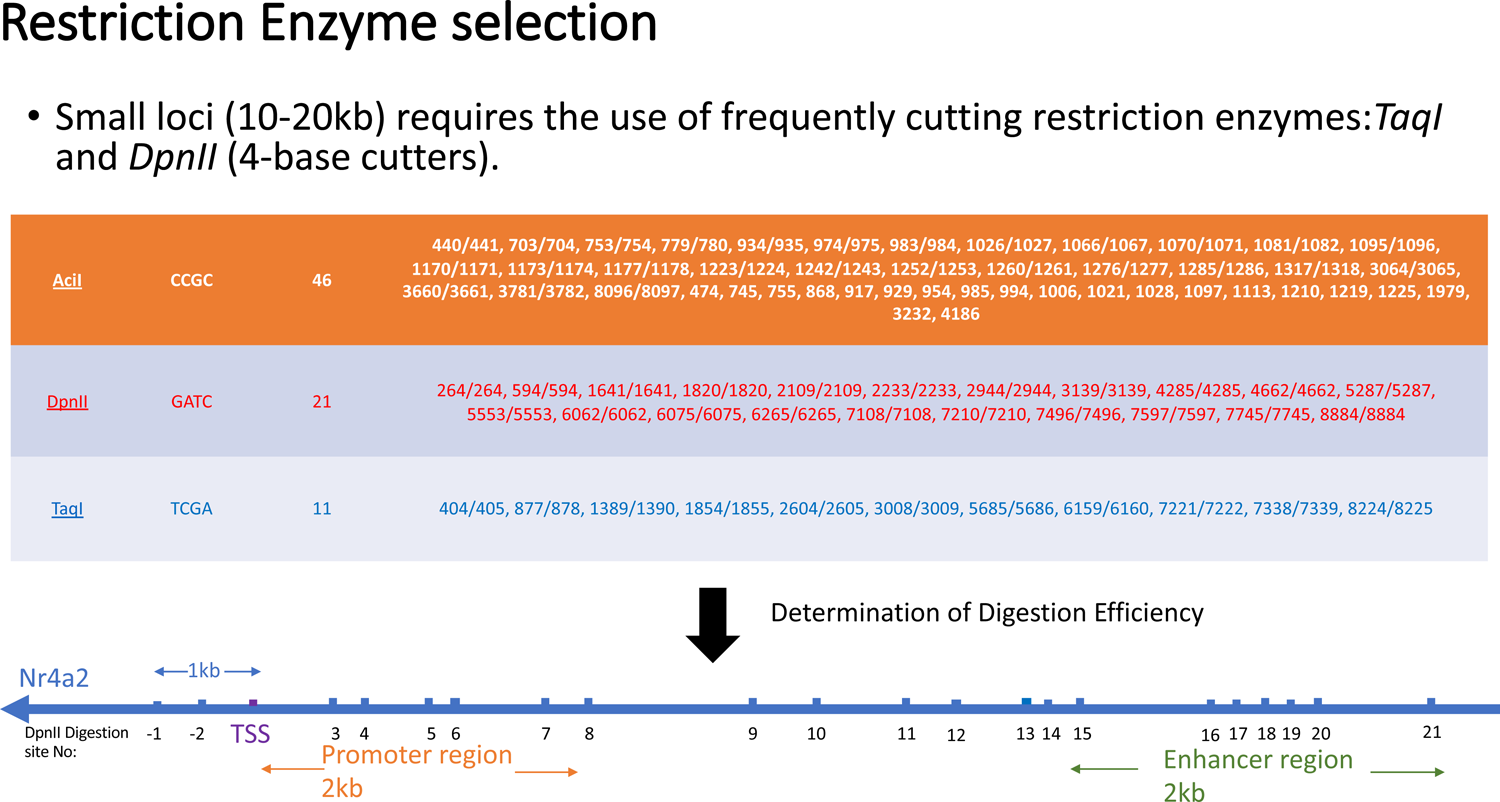
Potential ADRAM binding sites at the Nr4a2 promoter region. There are two sites located at −469∼496bp and −871∼-894bp upstream of TSS, with CREB binding sites in close proximity to both.

**Supplemental Figure 10.**
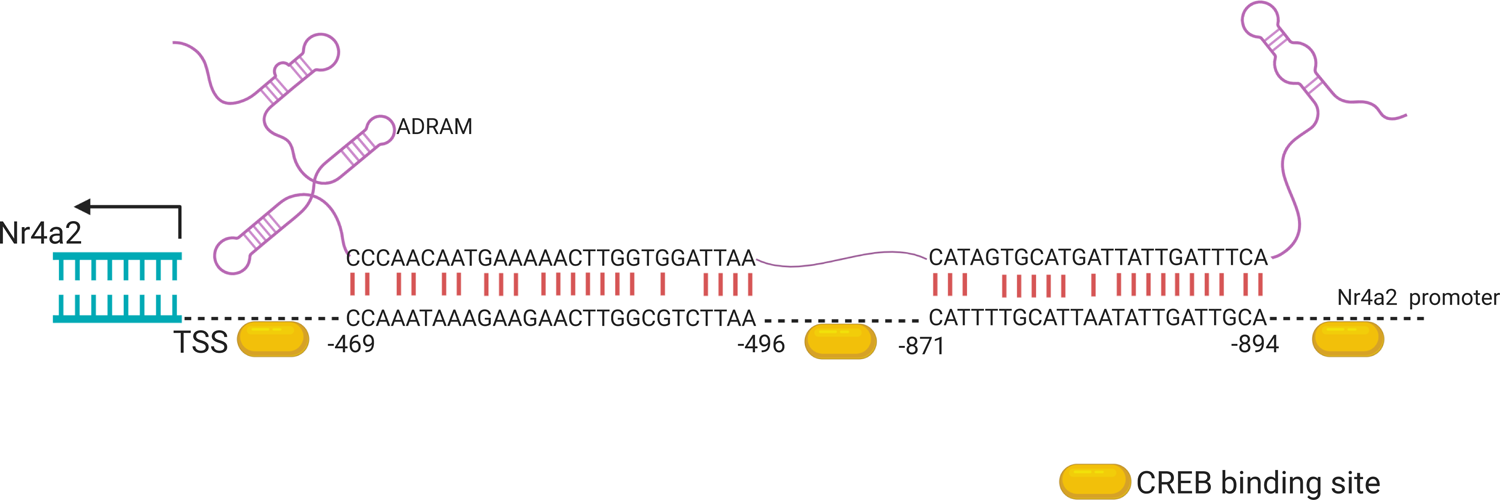
ADARM ASO-mediated knockdown inhibits extinction learning induced H3K4^me3^ occupancy at Nr4a2 promoter. chIP-qPCR result shows administration of ADRAM ASO inhibits H3K4me3 occupancy at the Nr4a2 promoter region at ILPFC post fear extinction training.

**Supplemental Figure 11.**
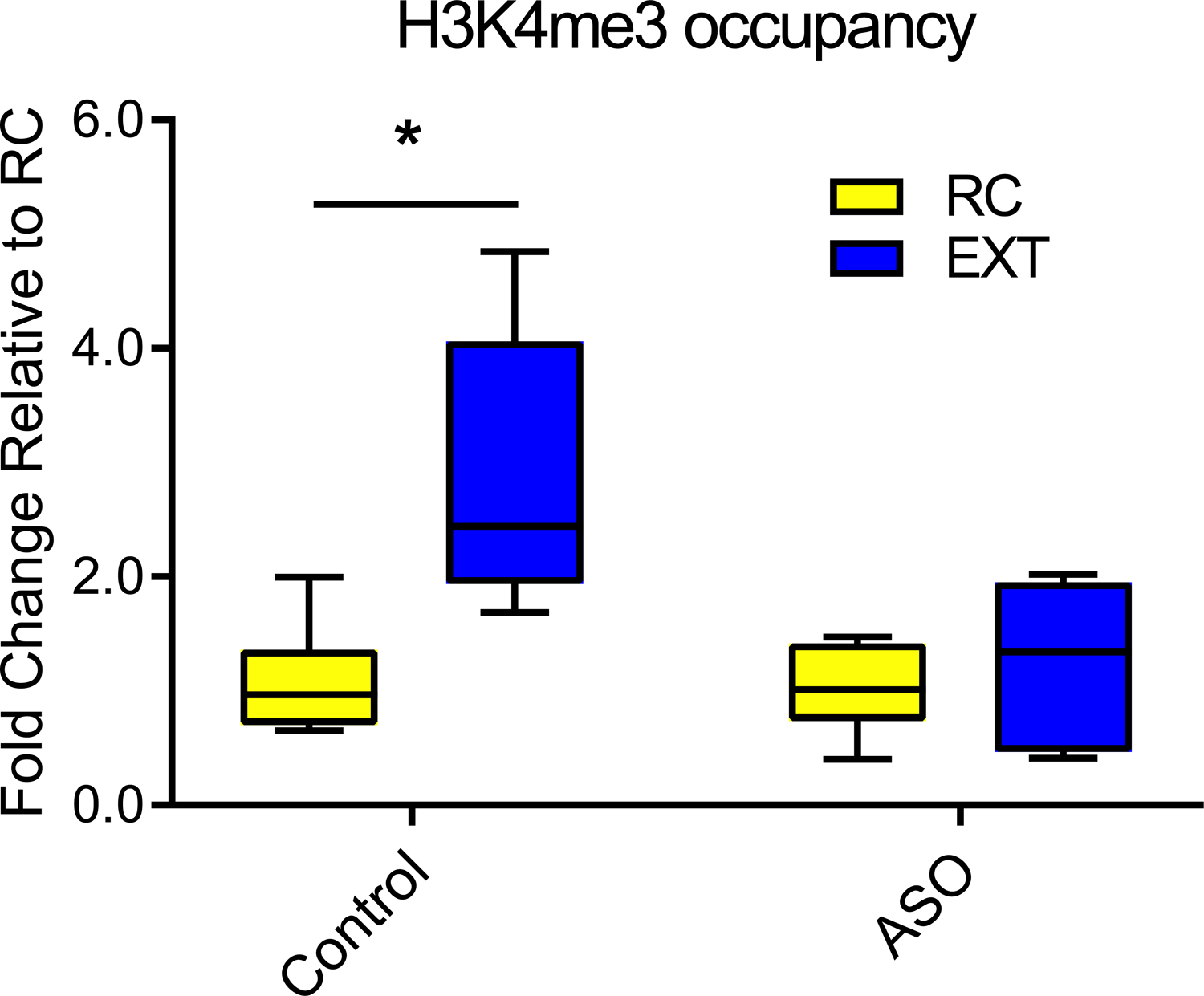
Representative silver stain of crosslinked ADRAM:protein complex derived from the ILPFC of EXT trained mice. M: marker, ADRAM: ADRAM probe enriched proteins and Con: Scrambled control probe enriched proteins.

**Supplemental Figure 12.**
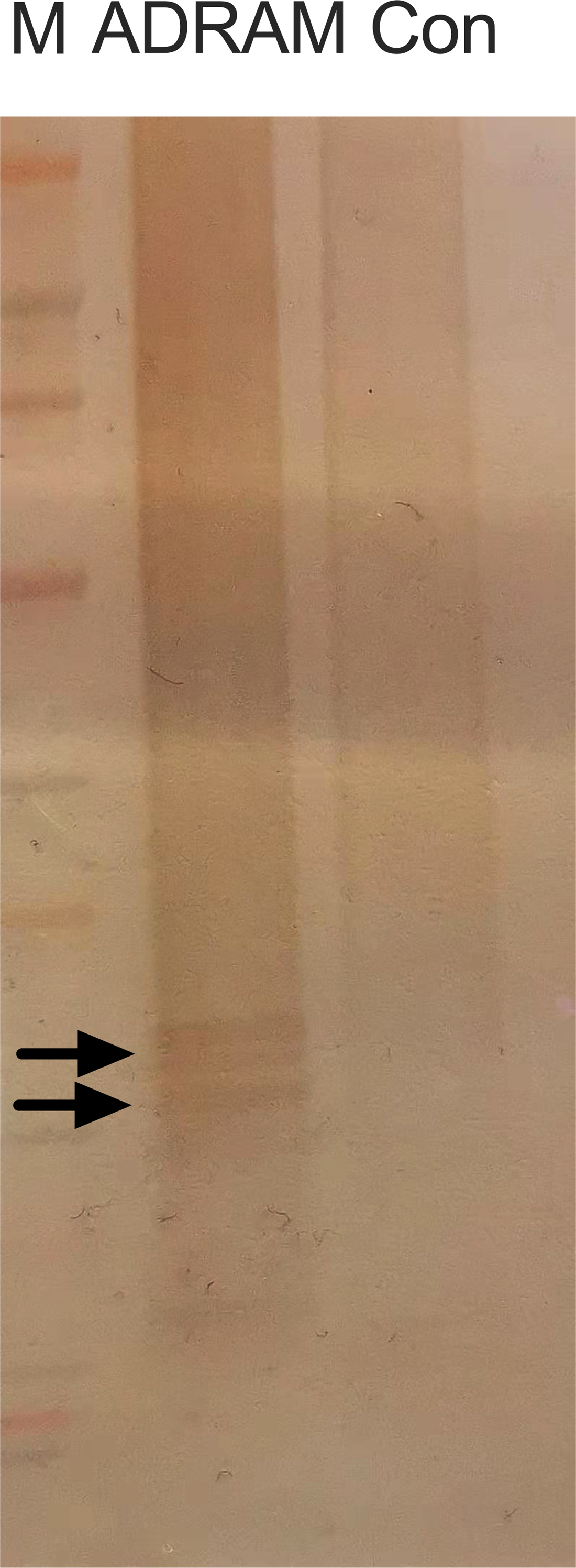
ADRAM is expressed in various regions of the brain. RT**-**qPCR reveals the expression of ADRAM in the cerebellum, prefrontal cortex, hippocampus, and somatosensory cortex.

**Supplemental Figure 13.**
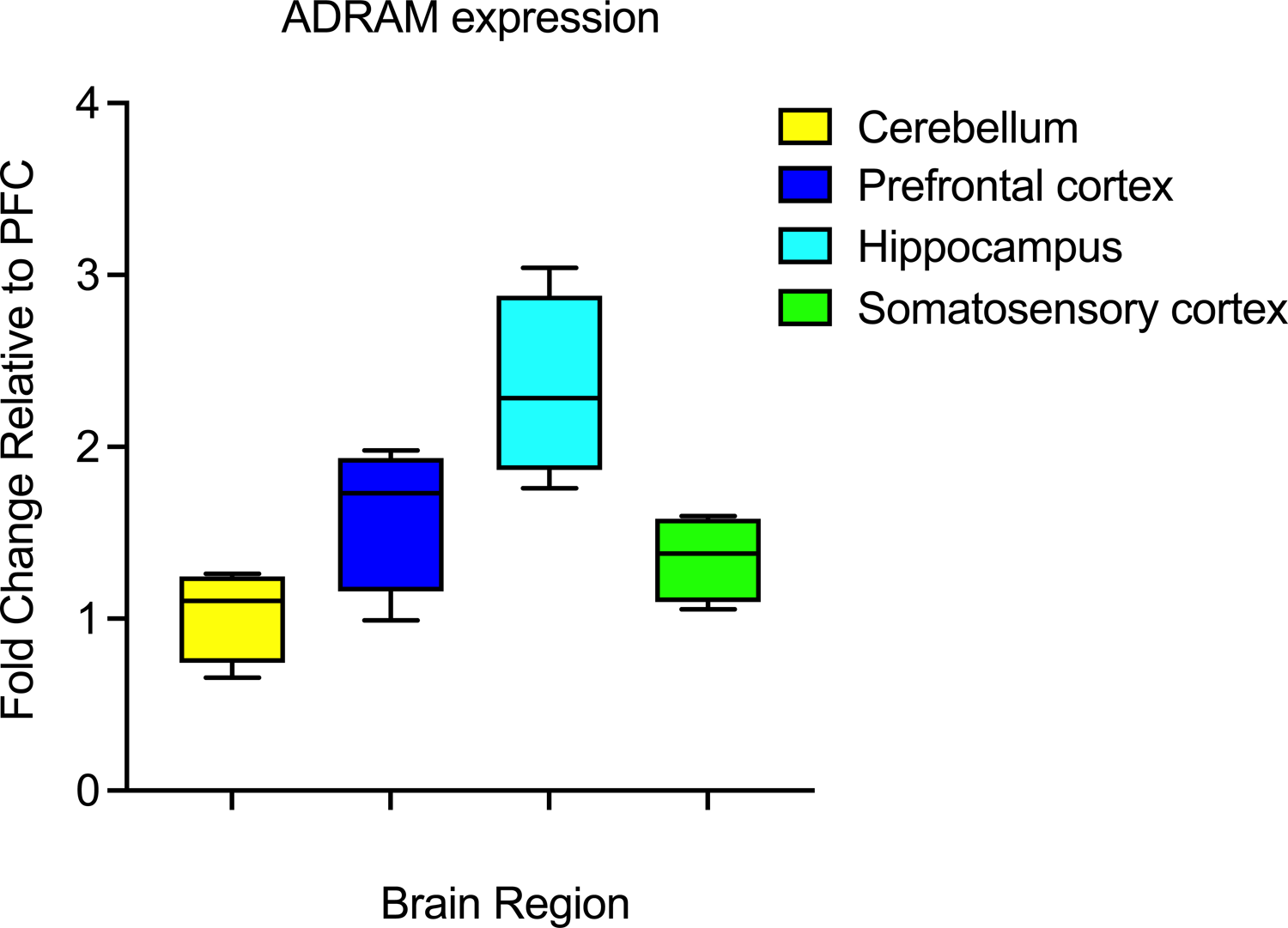
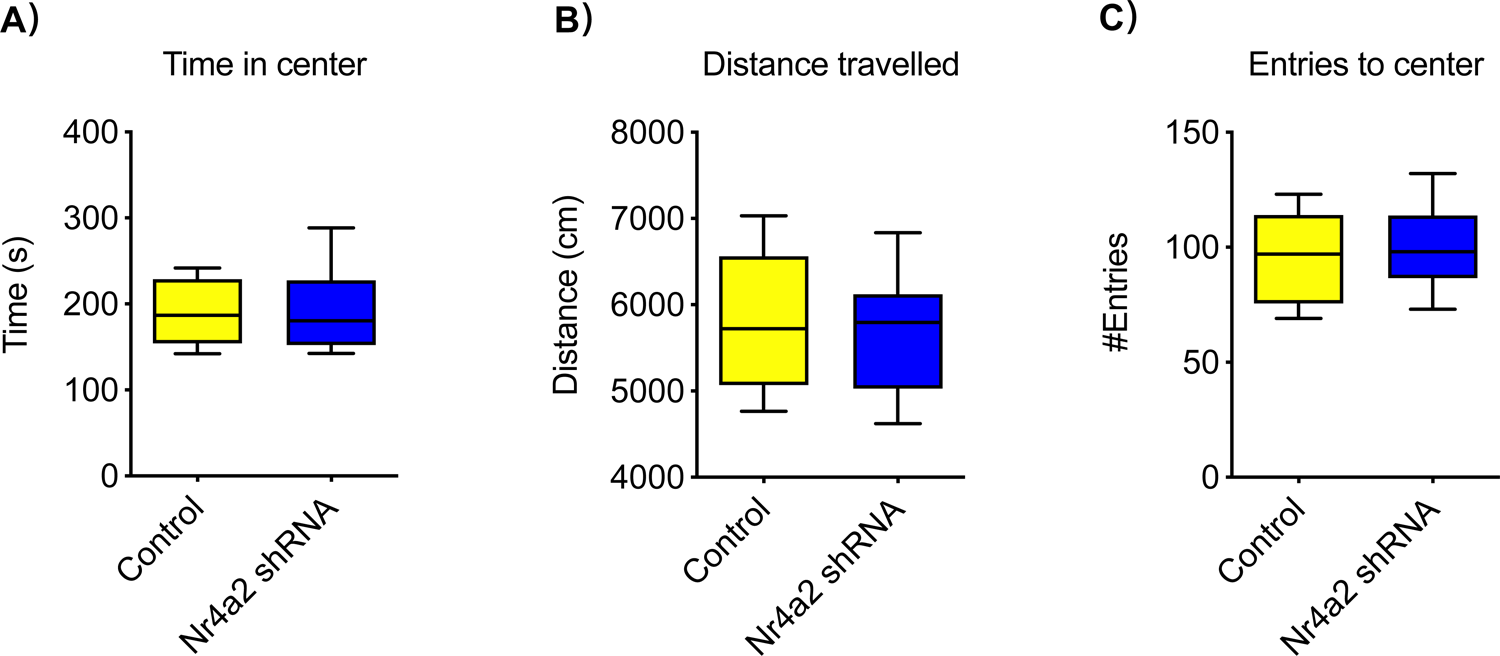
Nr4a2 knockdown has no effect on the anxiety-like behaviour. Compared with control virus treated mice, there is no effect of Nr4a2 shRNA-mediated knockdown on time spent in center A), distance travelled B) and entries to center C) (n=8-10 mice per group)

## REFERENCES

1. Alberini CM. (2009). Transcription factors in long-term memory and synaptic plasticity. Physiol Rev. 89:121–145.

2. Arab K, Karaulanov E, Musheev M, Trnka P, Schafer A, Grummt I, et al. (2019): GADD45A binds R-loops and recruits TET1 to CpG island promoters. Nat Genet 51:217–223.

3. Barry G, Briggs JA, Vanichkina DP, Poth EM, Beveridge NJ, Ratnu VS, et al. (2014): The long non-coding RNA Gomafu is acutely regulated in response to neuronal activation and involved in schizophrenia-associated alternative splicing. Mol Psychiatry 19:486–494.

4. Berg D, Holzmann C, Riess O (2003). 14-3-3 proteins in the nervous system. Nat Rev Neurosci 4:752–762.

5. Bose DA, Berger SL (2017). eRNA binding produces tailored CBP activity profiles to regulate gene expression. RNA Biol 14:1655–1659.

6. Butler AA, Johnston DR, Kaur S, Lubin FD (2019). Long noncoding RNA NEAT1 mediates neuronal histone methylation and age-related memory impairment. Sci Signal 12.

7. Bredy TW, Wu H, Crego C, Zellhoefer J, Sun YE, Barad M (2007). Histone modifications around individual BDNF gene promoters in prefrontal cortex are associated with extinction of conditioned fear. Learn Mem 14:268–276.

8. Bridi MS, Hawk JD, Chatterjee S, Safe S, Abel T (2017). Pharmacological Activators of the NR4A Nuclear Receptors Enhance LTP in a CREB/CBP-Dependent Manner. Neuropsychopharmacology 42:1243–1253.

9. Bruel-Jungerman E, Davis S, Laroche S. (2007). Brain plasticity mechanisms and memory: a party of four. Neuroscientist. 13:492–505.

10. Bussotti G, Leonardi T, Clark MB, Mercer TR, Crawford J, Malquori L, Notredame C, Dinger ME, Mattick JS, Enright AJ. (2016). Improved definition of the mouse transcriptome via targeted RNA sequencing.Genome Res. 26(5):705–16.

11. Cabili MN, Dunagin MC, McClanahan PD, Biaesch A, Padovan-Merhar O, Regev A, Rinn JL, Raj A. (2015). Localization and abundance analysis of human lncRNAs at single-cell and single-molecule resolution. Genome Biol. 16(1):20.

12. Cajigas I et al. (2018). The Evf2 ultraconserved enhancer lncRNA functionally and spatially organizes megabase distant genes in the developing forebrain. Molecular Cell 71: 956–972.

13. Carrieri C, Cimatti L, Biagioli M, Beugnet A, Zucchelli S, Fedele S, et al. (2012). Long non-coding antisense RNA controls Uchl1 translation through an embedded SINEB2 repeat. Nature 491:454–457.

14. Carullo NVN, Phillips Iii RA, Simon RC, Soto SAR, Hinds JE, Salisbury AJ, et al. (2020). Enhancer RNAs predict enhancer-gene regulatory links and are critical for enhancer function in neuronal systems. Nucleic Acids Res 48:9550–9570.

15. Chanda K, Das S, Chakraborty J, Bucha S, Maitra A, Chatterjee R, et al. (2018). Altered levels of Long ncRNAs Meg3 and Neat1 in cell and animal models of huntington’s disease. RNA Biol 15:1348–1363.

16. Chatterjee S, Walsh EN, Yan AL, Giese KP, Safe S, Abel T. (2020). Pharmacological activation of Nr4a rescues age-associated memory decline. Neurobiol Aging. 85:140–144.

17. Chen LF, Lin YT, Gallegos DA, Hazlett MF, Gomez-Schiavon M, Yang MG, et al. (2019). Enhancer histone acetylation modulates transcriptional bursting dynamics of neuronal activity-inducible genes. Cell Rep 26:1174–1188 e1175.

18. Chu C, Zhang QC, da Rocha ST, Flynn RA, Bharadwaj M, Calabrese JM, et al. (2015). Systematic discovery of Xist RNA binding proteins. Cell 161:404-416.

19. Cloutier SC, Wang S, Ma WK, Al Husini N, Dhoondia Z, Ansari A, et al. (2016). Regulated formation of lncRNA-DNA hybrids enables faster transcriptional induction and environmental adaptation. Mol Cell 61:393–404.

20. Deveson IW et al. (2018). Universal alternative splicing of noncoding exons. Cell Systems 6: 245–255.

21. Deveson IW, Hardwick SA, Mercer TR and Mattick JS (2017). The dimensions, dynamics, and relevance of the mammalian noncoding transcriptome. Trends in Genetics 33: 464–478.

22. Fan S, et al. (2020). lncRNA CISAL inhibits BRCA1 transcription by forming a tertiary structure at its promoter iScience. 23(2):100835.

23. Feng J, Shao N, Szulwach KE, Vialou V, Huynh J, Zhong C, Le T, Ferguson D, Cahill ME, Li Y, Koo JW, Ribeiro E, Labonte B, Laitman BM, Estey D, Stockman V, Kennedy P, Couroussé T, Mensah I, Turecki G, Faull KF, Ming GL, Song H, Fan G, Casaccia P, Shen L, Jin P, Nestler EJ. (2015). Role of Tet1 and 5-hydroxymethylcytosine in cocaine action. Nat Neurosci. 18(4):536–44

24. Foote M, Qiao H, Graham K, Wu Y, Zhou Y (2015). Inhibition of 14-3-3 proteins leads to schizophrenia-related behavioral phenotypes and synaptic defects in .ice. Biol Psychiatry 78:386–395.

25. Frazee, A.C. et al. (2015). Ballgown bridges the gap between transcriptome assembly and expression analysis. Nat. Biotechnol. 33, 243–246

26. Fullwood MJ, Ruan Y (2009): ChIP-based methods for the identification of long-range chromatin interactions. J Cell Biochem 107:30–39.

27. Giles N, Forrest A, Gabrielli B (2003). 14-3-3 acts as an intramolecular bridge to regulate cdc25B localization and activity. J Biol Chem 278:28580–28587.

28. Gräff J, Joseph NF, Horn ME, Samiei A, Meng J, Seo J, Rei D, Bero AW, Phan TX, Wagner F, Holson E, Xu J, Sun J, Neve RL, Mach RH, Haggarty SJ, Tsai LH.(2014). Epigenetic priming of memory updating during reconsolidation to attenuate remote fear memories. Cell. 2014 Jan 16;156(1-2):261-76.

29. Grinman E et al. (2021). Activity-regulated synaptic targeting of lncRNA ADEPTR mediates structural plasticity by localizing Sptn1 and AnkB in dendrites. Science Advances 7: eabf0605.

30. Hagege H, Klous P, Braem C, Splinter E, Dekker J, Cathala G, et al. (2007). Quantitative analysis of chromosome conformation capture assays (3C-qPCR). Nat Protoc 2:1722–1733.

31. Halder R, et al. (2016). DNA methylation changes in plasticity genes accompany the formation and maintenance of memory. Nat Neurosci. 19(1):102–10.

32. Hollensen AK et al. (2020). circZNF827 nucleates a transcription inhibitory complex to balance neuronal differentiation. eLife 9: e58478.

33. Issler O, van der Zee YY, Ramakrishnan A, Wang J, Tan C, Loh YE, et al. (2020). Sex-specific role for the long non-coding RNA LINC00473 in depression. Neuron 106:912–926 e915.

34. Jeon Y, Lee JT (2011). YY1 tethers Xist RNA to the inactive X nucleation center. Cell 146:119–133.

35. Kim TK, Hemberg M, Gray JM, Costa AM, Bear DM, Wu J, et al. (2010). Widespread transcription at neuronal activity-regulated enhancers. Nature 465:182–187.

36. Kim TK, Hemberg M, Gray JM (2015). Enhancer RNAs: a class of long noncoding RNAs synthesized at enhancers. Cold Spring Harb Perspect Biol 7:a018622.

37. Kim D, Langmead B, Salzberg SL (2015). HISAT: a fast spliced aligner with low memory requirements. Nat Methods 12:357–360.

38. Kukharsky MS, et al. (2020). Long non-coding RNA Neat1 regulates adaptive behavioural response to stress in mice. Transl Psychiatry, 10(1):171.

39. Labonte B, Abdallah K, Maussion G, Yerko V, Yang J, Bittar T, et al. (2020). Regulation of impulsive and aggressive behaviours by a novel lncRNA. Mol Psychiatry. 6:10.1038/s41380-019-0637-4.

40. Lepack AE, Werner CT, Stewart AF, Fulton SL, Zhong P, Farrelly LA, Smith ACW, Ramakrishnan A, Lyu Y, Bastle RM, Martin JA, Mitra S, O’Connor RM, Wang ZJ, Molina H, Turecki G, Shen L, Yan Z, Calipari ES, Dietz DM, Kenny PJ, Maze I. (2020). Dopaminylation of histone H3 in ventral tegmental area regulates cocaine seeking. Science. 368(6487):197–201.

41. Li D, Zhang J, Wang M, Li X, Gong H, Tang H, Chen L, Wan L, Liu Q. (2018). Activity dependent LoNA regulates translation by coordinating rRNA transcription and methylation.Nat Commun. 2018 Apr 30;9(1):1726.

42. Li X, Wei W, Zhao QY, Widagdo J, Baker-Andresen D, Flavell CR, et al. (2014). Neocortical Tet3-mediated accumulation of 5-hydroxymethylcytosine promotes rapid behavioral adaptation. Proc Natl Acad Sci U S A 111:7120–7125.

43. Li X, Zhao Q, Wei W, Lin Q, Magnan C, Emami MR, et al. (2019). The DNA modification N6-methyl-2’-deoxyadenosine (m6dA) drives activity-induced gene expression and is required for fear extinction. Nat Neurosci 22:534–544.

44. Li H, Durbin R (2009). Fast and accurate short read alignment with Burrows-Wheeler transform. Bioinformatics 25:1754–1760.

45. Li H, Handsaker B, Wysoker A, Fennell T, Ruan J, Homer N, Marth G, Abecasis G, Durbin R, and 1000 Genome Project Data Processing Subgroup (2009). The Sequence alignment/map (SAM) format and SAMtools, Bioinformatics 25(16) 2078–9

46. Li Y, Syed J, Sugiyama H (2016). RNA-DNA triplex formation by long noncoding RNAs. Cell Chem Biol 23:1325–1333.

47. Liau WS, Samaddar S, Banerjee S, Bredy TW (2021). On the functional relevance of spatiotemporally-specific patterns of experience-dependent long noncoding RNA expression in the brain. RNA Biol:1–12.

48. Liu N, Zhou KI, Parisien M, Dai Q, Diatchenko L, Pan T. (2017). N6-methyladenosine alters RNA structure to regulate binding of a low-complexity protein. Nucleic Acids Res. 45(10):6051–6063.

49. Ma M, Xiong W, Hu F, Deng MF, Huang X, Chen JG, Man HY, Lu Y, Liu D, Zhu LQ. (2020). A novel pathway regulates social hierarchy via lncRNA AtLAS and postsynaptic synapsin IIb. Cell Res. 30(2):105–118.

50. Malik AN, Vierbuchen T, Hemberg M, Rubin AA, Ling E, Couch CH, et al. (2014). Genome-wide identification and characterization of functional neuronal activity-dependent enhancers. Nat Neurosci 17:1330–1339.

51. Martin H, Rostas J, Patel Y, Aitken A (1994). Subcellular localisation of 14-3-3 isoforms in rat brain using specific antibodies. J Neurochem 63:2259–2265.

52. Martin SJ, Grimwood PD, Morris RG. (2000). Synaptic plasticity and memory: an evaluation of the hypothesis. Annu Rev Neurosci. 23:649–711.

53. Marzinke MA, Mavencamp T, Duratinsky J, Clagett-Dame M (2013). 14-3-3epsilon and NAV2 interact to regulate neurite outgrowth and axon elongation. Arch Biochem Biophys 540:94–100.

54. McKinsey T.A., Zhang C.L., Lu J., Olson E.N. (2000). Signal-dependent nuclear export of a histone deacetylase regulates muscle differentiation. Nature. 408:106–111.

55. McNulty SE, Barrett RM, Vogel-Ciernia A, Malvaez M, Hernandez N, Davatolhagh MF, Matheos DP, Schiffman A, Wood MA (2012). Differential roles for Nr4a1 and Nr4a2 in object location vs. object recognition long-term memory.Learn Mem. 19(12):588–92.

56. McQuown SC, Barrett RM, Matheos DP, Post RJ, Rogge GA, Alenghat T, Mullican SE, Jones S, Rusche JR, Lazar MA, Wood MA (2011). HDAC3 is a critical negative regulator of long-term memory formation. Neurosci. 31(2):764–74.

57. Mercer TR, Dinger ME, Sunkin SM, Mehler MF, Mattick JS (2008). Specific expression of long noncoding RNAs in the mouse brain. Proc Natl Acad Sci U S A 105:716–721.

58. Mercer TR, Mattick JS. (2013). Structure and function of long noncoding RNAs in epigenetic regulation. Nat Struct Mol Biol. 20(3):300–7.

59. Mercer TR, Gerhardt DJ, Dinger ME, Crawford J, Trapnell C, Jeddeloh JA, Mattick JS, Rinn JL.(2011). Targeted RNA sequencing reveals the deep complexity of the human transcriptome. Nat Biotechnol. 30(1):99–104.

60. Mercer TR, Clark MB, Crawford J, Brunck ME, Gerhardt DJ, Taft RJ, Nielsen LK, Dinger ME, Mattick JS. (2014). Targeted sequencing for gene discovery and quantification using RNA CaptureSeq. Nat Protoc. 9(5):989–1009.

61. Pertea M, Kim D, Pertea GM, Leek JT, Salzberg SL (2016). Transcript-level expression analysis of RNA-seq experiments with HISAT, StringTie and Ballgown. Nat Protoc 11:1650–1667.

62. Pertea M, Pertea GM, Antonescu CM, Chang TC, Mendell JT & Salzberg SL. (2015). StringTie enables improved reconstruction of a transcriptome from RNA-seq reads Nature Biotechnology, doi:10.1038/nbt.3122

63. Perry RB-T, Hezroni H, Goldrich MJ and Ulitsky I (2018). Regulation of neuroregeneration by long noncoding RNAs. Molecular Cell 72: 553–567.

64. Qiao H, Foote M, Graham K, Wu Y, Zhou Y (2014). 14-3-3 proteins are required for hippocampal long-term potentiation and associative learning and memory. J Neurosci 34:4801–4808.

65. Quinlan AR, Hall IM. (2010). BEDTools: a flexible suite of utilities for comparing genomic features. Bioinformatics. 26(6):841–2.

66. Raveendra BL et al. (2018). Long noncoding RNA GM12371 acts as a transcriptional regulator of synapse function. Proceedings of the National Academy of Sciences USA 115: E10197–E10205.

67. Sando R3rd, Gounko N, Pieraut S, Liao L, Yates J3rd, Maximov A. (2012). HDAC4 governs a transcriptional program essential for synaptic plasticity and memory. Cell. 151(4):821–834.

68. Seiler J, Breinig M, Caudron-Herger M, Polycarpou-Schwarz M, Boutros M, Diederichs S. (2017). The lncRNA VELUCT strongly regulates viability of lung cancer cells despite its extremely low abundance. Nucleic Acids Res. 45(9):5458–5469.

69. Spadaro PA, Bredy TW. (2012). Emerging role of non-coding RNA in neural plasticity, cognitive function, and neuropsychiatric disorders. Front Genet. 3:132.

70. Spadaro PA, Flavell CR, Widagdo J, Ratnu VS, Troup M, Ragan C, et al. (2015). Long noncoding RNA-directed epigenetic regulation of gene expression is associated with anxiety-like behavior in mice. Biol Psychiatry 78:848–859.

71. Tang WH, Shilov IV, Seymour SL. (2008). Nonlinear fitting method for determining local false discovery rates from decoy database searches. Journal of Proteome Research. 7:3661–3667.

72. Vecsey CG, Hawk JD, Lattal KM, Stein JM, Fabian SA, Attner MA, Cabrera SM, McDonough CB, Brindle PK, Abel T, Wood MA.(2007). Histone deacetylase inhibitors enhance memory and synaptic plasticity via CREB:CBP-dependent transcriptional activation. J Neurosci. 27(23):6128–40.

73. Wakeling E, et al. (2021). Missense substitutions at a conserved 14-3-3 binding site in HDAC4 cause a novel intellectual disability syndrome. HGG Adv. 2(1):100015.

74. Wang X, Cairns MJ, Yan J (2019). Super-enhancers in transcriptional regulation and genome organization. Nucleic Acids Res 47:11481–11496.

75. Wang F et al. (2021). The long noncoding RNA Synage regulates synapse stability and neuronal function in the cerebellum. Cell Death & Differentiation in press.

76. Wei W, Coelho CM, Li X, Marek R, Yan S, Anderson S, et al. (2012). p300/CBP-associated factor selectively regulates the extinction of conditioned fear. J Neurosci 32:11930–11941.

77. Wu Q, Yi X (2018). Down-regulation of Long Noncoding RNA MALAT1 Protects Hippocampal Neurons Against Excessive Autophagy and Apoptosis via the PI3K/Akt Signaling Pathway in Rats with Epilepsy. J Mol Neurosci 65:234–245.

78. Xiong Z, Lo HP, McMahon KA, Martel N, Jones A, Hill MM, Parton RG, Hall TE. (2021). In vivo proteomic mapping through GFP-directed proximity-dependent biotin labelling in zebrafish. Elife. 10:e64631.

79. Xu H, Brown AN, Waddell NJ, Liu X, Kaplan GJ, Chitaman JM, et al. (2020). Role of long noncoding RNA Gas5 in cocaine action. Biol Psychiatry 88:758–766.

80. Yamazaki T, Souquere S, Chujo T, Kobelke S, Chong YS, Fox AH, et al. (2018). Functional domains of NEAT1 architectural lncRNA induce paraspeckle assembly through phase separation. Mol Cell 70:1038–1053 e1037.

81. Zhang Y, Liu T, Meyer CA, Eeckhoute J, Johnson DS, Bernstein BE, et al. (2008). Model-based analysis of ChIP-Seq (MACS). Genome Biol 9:R137.

82. Zhang J, Zhou Y (2018). 14-3-3 proteins in glutamatergic synapses. Neural Plast 2018:8407609.

83. Zhu Y, Huang M, Bushong E, Phan S, Uytiepo M, Beutter E, Boemer D, Tsui K, Ellisman M, Maximov A. (2019). Class IIa HDACs regulate learning and memory through dynamic experience-dependent repression of transcription. Nat Commun. 10(1):3469.

